# Persistently increased post-stress activity of paraventricular thalamic neurons is essential for the emergence of stress-induced maladaptive behavior

**DOI:** 10.1101/2023.04.20.537629

**Authors:** Anna Jász, László Biró, Zsolt Buday, Bálint Király, Orsolya Szalárdy, Krisztina Horváth, Gergely Komlósi, Róbert Bódizs, Krisztina J. Kovács, Marco A. Diana, Balázs Hangya, László Acsády

## Abstract

**ABSTRACT:** Traumatic events can immediately lead to debilitating symptoms collectively called Acute Stress Disorder (ASD), however the mechanisms of ASD are poorly understood. Using a rodent model of ASD here we identify a crucial communication bottleneck between the brainstem and the forebrain, the calretinin-positive neurons in the paraventricular thalamus (PVT/CR+), that controls ASD. We show that following a single acute stress event, the pre-sleep behavior of the mice is altered for several days in parallel with a persistent increase in the firing rate of PVT/CR+ cells. Optogenetic inhibition of PVT/CR+ neuronal activity after the stress event for one hour was sufficient to rescue both the ASD symptoms and the prolonged increase in PVT/CR+ firing rate. Inhibition applied 5 days later was still able to ameliorate some of the symptoms. These data suggest that post-stress activity of PVT/CR+ neurons play a critical role in mediating the acute forms of stress-related affective dysfunctions.

**One-Sentence Summary:** Post-stress inhibition of paraventricular thalamic neurons prevents the emergence of acute stressed phenotype.

---

Acute stress disorder (ASD) can develop following various types of traumas including verbal or physical abuse, serious injury, and threat of sudden death of a loved one^1, 2^. If ASD lasts longer than 30 days, it is called posttraumatic stress disorder (PTSD).^1, 2^ Similar to PTSD the main symptoms of ASD in humans include intense psychological distress, hypervigilance, and sleep disturbances^2^ . Despite the high prevalence of ASD, the majority of stress research focused on chronic stress and PTSD and few studies have focused on the peri-trauma time window^3–5^. As a consequence, the effects of an intense stress event on post-stress neuronal activity in parallel with stress-induced behavioral alterations have not been intensively studied^5^. Because the treatment of ASD is significantly more effective than that of PTSD^1^ and 70% of the patients diagnosed with ASD develop PTSD^1^ understanding ASD mechanisms is highly relevant. In this project, we used a rodent ASD model and aimed to understand the neuronal basis of behavioral alterations following an intense stress event.

Calretinin-positive neurons of the paraventricular thalamus (PVT/CR+) represent a critical bottleneck in the communications between the brainstem and forebrain systems which controls arousal and the behavioral response to aversive stimuli.^6, 7^ PVT/CR+ cells receive highly selective inputs from a variety of hypothalamic and brainstem centers^7, 8^ which transfer salience^9^, valance^10^, arousal^7, 11, 12^, satiety^13–15^, social^16^, stress ^17^and fear^18, 19^ signals via the PVT^8, 20, 21^ to the forebrain. The output of PVT/CR+ cells is also unique^7, 22^ . These cells represent the only subcortical, glutamatergic cell population that innervate all major forebrain centers (including, amygdala, medial prefrontal cortex, hippocampal formation, and n. accumbens)^7^ involved in stress and fear related behavioral responses. Accordingly, PVT has been implicated in highly diverse affective and metabolic conditions like obesity^14, 23, 24^ and addiction.^25, 26^ The organization and inputs of PVT/CR+ neurons are highly conserved between mice and humans^7^.in humans, similar to rodents, both lesion and functional imaging studies link PVT to altered vigilance states.^27–29^

Recent data indicate that PVT is involved in stress responses as well^17, 21, 30–34^ and PVT cells can display a transient increase in activity when exposed to various stressors^30, 35^. Whether this increase in activity reflects a transient elevation due to increased arousal^7, 12^ or persist for longer duration and PVT/CR+ cells can potentially have a specific role in establishing ASD is presently unclear. Thus, in this study, using a well-established stress model based on innate fear^36–39^ we tested the role of PVT/CR+ cells in ASD. We found that the activity PVT/CR+ neurons display prolonged changes following a stress event and the elevated post-stress activity of PVT/CR+ neurons is causally involved in the manifestation of ASD-like behavior in mice.

## RESULTS

### A rodent model of ASD, optogenetic inhibition of PVT/CR+ cells

In order to test the role of PVT/CR+ neurons in establishing the acute stressed phenotype we used a behavioral essay focusing on the peri-trauma time window. Unescapable, predator odor stress exposure (POSE) is an efficient and well-accepted model for stress-induced long-term alteration in behavior^37, 38^. Accordingly, after recording the pre-stress behavior in the home cage (PRE 1-5 days) we exposed mice to 2-methyl-2-thiazoline (2MT) for 10 min^40, 41^ in a novel cage. We tested the causative role of PVT/CR+ cells in establishing the stressed phenotype by precisely timed optogenetic interference with their activity for one hour (2 sec ON 13 sec OFF duty cycle) after the stress exposure, when the animals were returned to their home cage using the inhibitory opsin SwiChR in CR-Cre mice (Fig. 1A and fig. S1). We used tetrode recordings to demonstrate the effect of SwiChR activation. PVT/CR+ cells decreased their firing rate during the laser ON periods (fig. S1), as expected. Since we applied optogenetic inhibition after, not during, POSE we avoided interference with perceiving the stress exposure. Following the stress day (POST 0) we continued to monitor the behavior of the animals for four more days (POST 1-4 days) in the home cage and compared home cage observatoins in the pre and the post-stress periods (POST 1-4 days). As a control for the SwiChR group we used AAV-DIO-EYFP injected stressed animals (EYFP or non-inhibited group). As a control for the stress situation, we used no odor exposure (NOE) and home cage groups (see Materials and Methods) as well.

**Fig 1.**
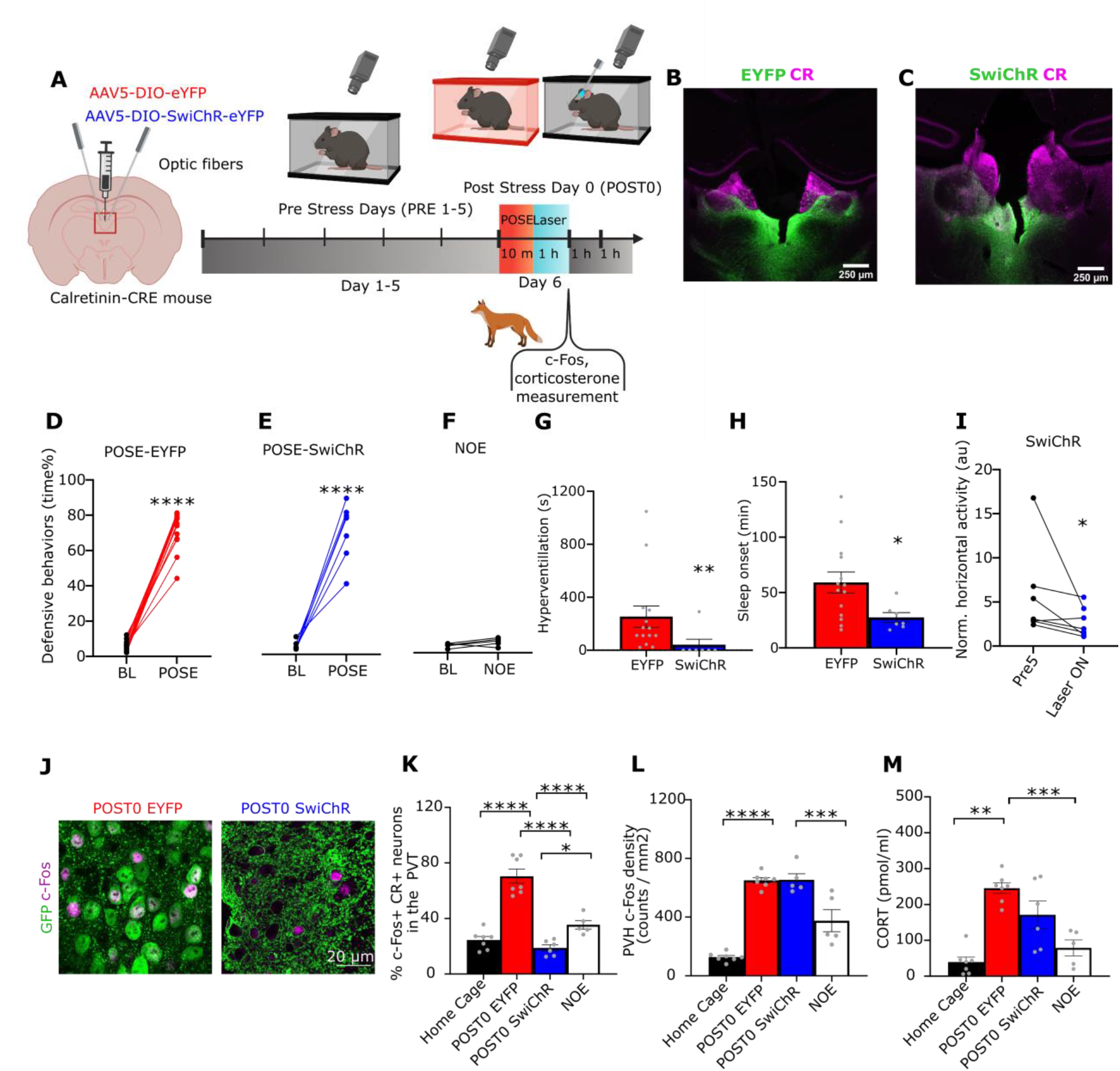
Short term effects of inhibiting PVT/CR+ neurons after the stress exposure. (A) Scheme of the experiment. AAVs were injected into PVT and behavior was assessed 4 weeks later. Following a 5 day baseline period to assess home cage behaviors (PRE5), mice were submitted to predator odor stress exposure (POSE). Immediately after the POSE both SwiChR and EYPF groups received a 1 hour long, photoinhibition of PVT/CR+ neurons in the home cage. (B-C) Representative confocal images showing EYFP and SwiChR expression in CR-positive (CR+) PVT neurons. Scale bars, 250 µm. (D-E) Quantification of time spent with defensive behavior during predator odor (2MT) stress exposure (POSE) in EYFP (D, t[13] = 21.05) and SwiChR (E, t[6] = 9.35) mice compared to a 2 min baseline (BL) period. (F) Same for the the no odor exposure (NOE, without 2MT presentation) group (t[4] = 1.822). (G-H) Bar graphs showing the duration of hyperventilation (G) and sleep onset latency (H) in EYFP (n=14, U = 10) and SwiChR (n=7, t[19] = 2.283) mice in their home cage after POSE. (I) Averaged, normalized horizontal activity during PVT/CR+ photoinhibition in the SwiChR animals (1^st^ hour of POST0 day) compared to the same period of the PRE 5 day (W = -26). (J) Confocal images showing c-Fos expression in PVT/CR+ neurons in EYFP and SwiChR mice one hour after POSE. Note that EYFP is a cytosolic labeling whereas SwichR is expressed in membranes. (K-L) Quantification of c-Fos expression in the PVT/CR+ cells (K, F(3,21) = 44.75, p < 0.0001) and in the paraventricular nucleus of the hypothalamus (PVH) (L) one hour after POSE in the four experimental groups; Home Cage control (n=7); EYFP (n=7); SwiChR (n=6); NOE no odor exposure control (F(3,20) = 50.00, p < 0.0001). (M) CORT level in the blood of the four experimental groups one hour after POSE (F(3,21) = 16.96, p < 0.0001). See Table S1 for the full results of the statistical tests. Data are shown as mean ± SEM. *p < 0.05, **p < 0.01, ***p < 0.01, ****p < 0.001

Since the SwiChR group was not inhibited during POSE both the EYFP and the SwiChR groups displayed robust defensive behaviors (escape jumps and freezing) as a response to 2MT exposure (Fig. 1, B, C D and E, fig. S2, A, Movie S1) in agreement with earlier data^42^. An increase in defensive behaviors was not observed in the NOE group (Fig. 1, F and Movie S2) showing that the novel environment per se did not contribute to the defensive behaviors. Following the return of the animals to their home cage, EYFP mice displayed significantly more high-frequency respiration (hyperventilation, Fig. 1G and Movie S3), and elevated sleep onset latency (Fig. 1H) on the day of POSE (POST0 day) compared to the SwiChR group (Fig. 1, H and I, Movie S4) which underwent 1 hour long photoinhibition after POSE. This indicates that inhibition of PVT/CR+ neurons after the stress can ameliorate the immediate impact of stress on behavioral symptoms. During the optogenetic inhibition the SwiChR animals did not display overt behavioral alterations, they showed normal locomotor activity. However, agreeing with a previous report^7^ they moved less compared to the identical period of their last pre-stress day (PRE5) (Fig. 1I and Movie S4).

One hour after POSE PVT/CR+ neurons in the EYFP animals displayed significantly increased c-Fos expression (Fig. 1, J and K) relative to home cage and NOE controls. In the EYFP mice, c-Fos expression also increased in the paraventricular hypothalamic nucleus (PVH) (Fig. 1L) together with significantly elevated blood corticosterone (CORT) levels (Fig. 1M) as in earlier studies^43, 44^. Photoinhibition of PVT/CR+ neurons for one hour after stress abolished the increase in c-Fos expression of PVT/CR+ cells (Fig. 1K). A decrease in c-Fos expression was most prominent under the fiber optic targeting PVT (fig. S2, B and C). Activation of SwiChR in PVT/CR+ cells, however, did not result in decreased c-Fos activity in PVH (Fig. 1L) or decreased blood corticosterone levels (Fig. 1M).

These short-term changes demonstrated that POSE could induce behavioral, hormonal, and gene expression alterations consistent with strong stress exposure^5, 45^. They also show that post stress photoinhibition of PVT/CR+ neurons is able to alleviate the immediate behavioral changes induced by POSE without influencing the activation of the HPA axis. This is in line with previous reports indicating that PVT is involved in the stress response, but the acute activation of the hypothalamic-pituitary-adrenal axis (HPA) is controlled by neuronal mechanisms other than PVT/CR+ neuronal activity^46, 47^.

### Prolonged effects of predator odor exposure on behavior

To test whether behavioral symptoms evoked by POSE persist for several days, mimicking human ASD, we compared the spontaneous wake and sleep behavior of the EYFP mice during the five pre (PRE1-5), and four post-stress days (POST1-4) outside and inside the nest (Fig. 2A). In order to focus on prolonged behavioral changes, the day of the stress (POST0 day), was not included in the analysis. Compared to PRE1-5 days, the horizontal locomotor activity of the EYFP animals remained significantly elevated for the entire duration of the POST period (POST1-4 days) (Fig. 2, B, C, D and E). After the stress, the animals frequently moved along the edge of their cages (Fig. 2I). After the stress event, the animals needed significantly more time to fall asleep (time between entering the nest and falling asleep, referred to as nest time, Fig. 2F), displayed significantly less nest-building behavior (Fig. 2G and Movie S5) but increased the time spent with freezing in the nest (Fig. 2H and Movie S6). We measured the quality of sleep by the frequency characteristics (spectral slope) of the EEG (see Methods)^48^. On the stress day (POST0) the spectral slope of the EEG was significantly increased during NREM sleep (fig. S2, D) consistent with the earlier data^49, 50^. This value, however, did not remain significant during the rest of the post-stress days (POST1-4) indicating normal sleep depth during this period (Fig. 2I).

**Fig 2.**
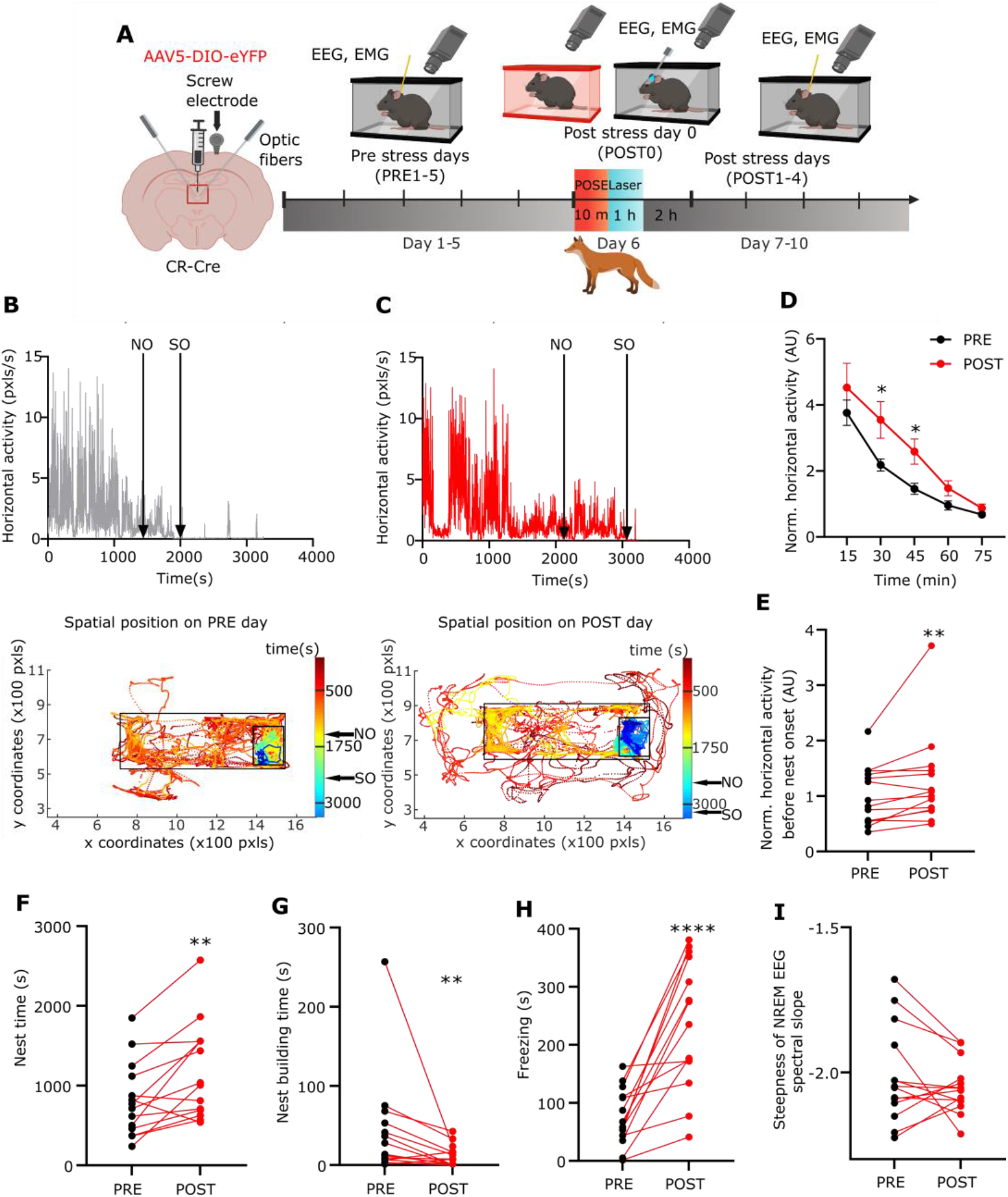
Long-term behavioral alterations after stress exposure. (A) Scheme of the experiment. (B-C) *Top,* Representative horizontal locomotor activity during a 3 hour session in one EYFP mouse within the home cage during one of the PRE days. Nest onset (NO) and sleep onset (SO) are indicated with black arrows. *Bottom,* Tracked movements of the same animal in the first hour. The dots show the spatial position of the head of the mouse in every frame. Color of the dots indicate time elapsed (s), from red to blue; small black rectangle marks the nest area; big black rectangle marks cage area. NO and SO, black arrows. (C) Same as H during a POST day. Head positions outside the cage indicate events when the animal was moving along the edge of its cage. (D) Temporal dynamics of averaged, normalized horizontal locomotor activity of EYFP the animals (n=14) during the PRE1-5 days (black) vs the POST1-4 days (red) periods (30 min, t[13] = 2.438; 45 min, t[13] = 2.392). (E) The averaged, normalized horizontal locomotor activity of EYFP the animals before sleep onset (W = 85). Dots represent the averaged daily values of individual animals. (F-I) (F) Nest time (t[13] = 3.579), (G) nest building time (W = -89), (H) freezing time (t[13] = 5.692) and (I) steepness of the NREM EEG spectral slope (t[13] = 1.086) in EYFP animals (n=14) during the PRE (black dots) and POST (red dots) period. Dots represent the averaged daily values of individual animals. See Table S1 for the full results of the statistical tests. Data are shown as mean ± SEM . *p < 0.05, **p < 0.01, ***p < 0.01, ****p < 0.001.

These data together demonstrate long-term, stress-induced changes in the spontaneous behavior of mice similar to human ASD symptoms including hypervigilance, difficulty in falling asleep, and intense emotional distress.

### Early alterations of PVT/CR+ neuronal activity after stress exposure

Next, we aimed to identify how the activity of PVT/CR+ neurons is altered in the post stress period relative to the pre-stress period. Earlier data indicated decreased GABA-A receptor mediated inhibition in PVT neurons following restraint stress^31^. Thus, we first tested functional GABA-A receptor-mediated synaptic currents after POSE in PVT/CR+ cells using *in vitro* slice preparations 2 hours after POSE (Fig. 3A) and compared them to a NOE group. We found that the frequency of spontaneous inhibitory synaptic currents (sIPSCs) significantly decreased in PVT/CR+ neurons (Fig. 3, B and C), while their amplitude remained unaltered (Fig. 3D). These data show that the PVT/CR+ cells receive decreased inhibitory synaptic activity immediately after the stress.

**Fig. 3.**
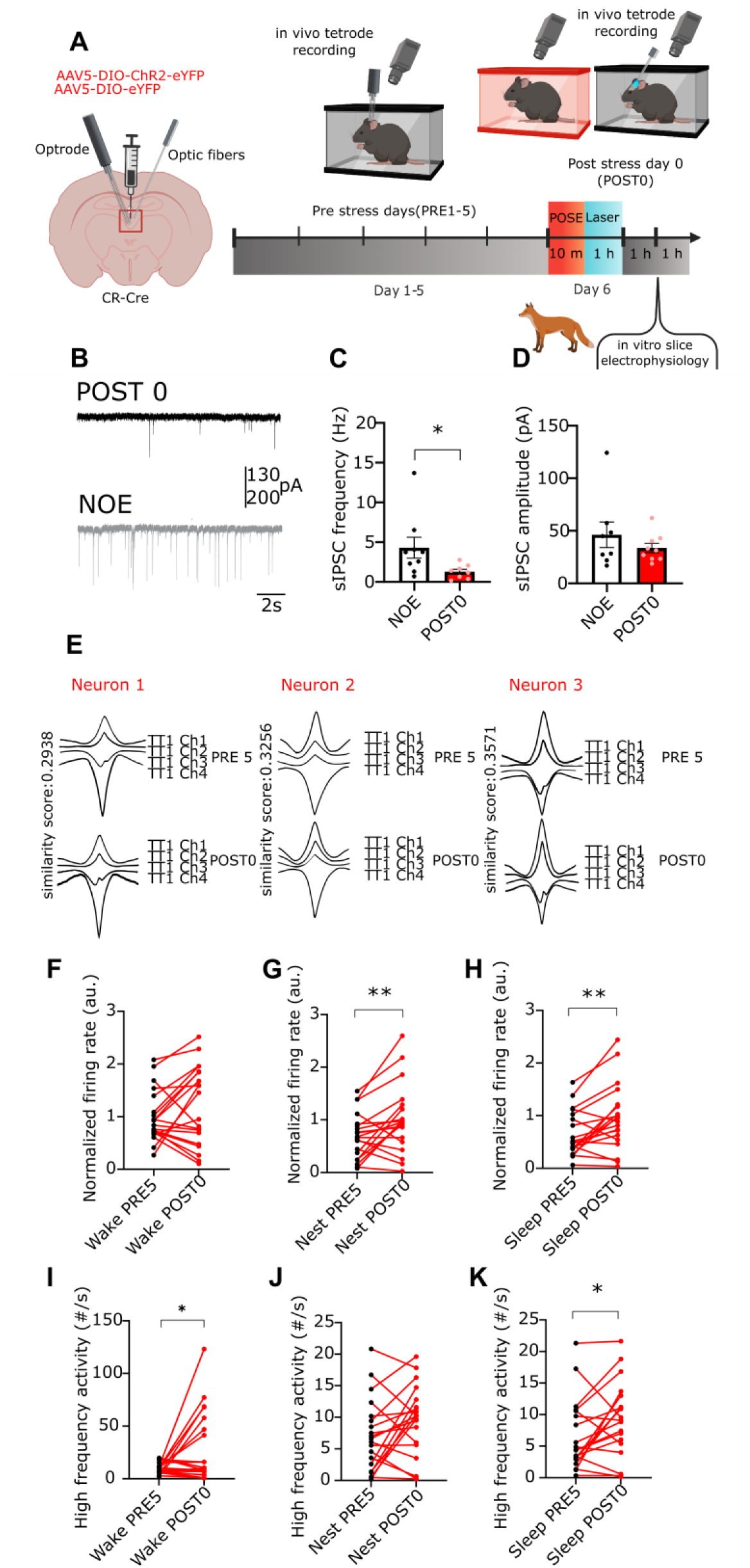
Short-term changes in PVT/CR+ unit activity after predator odor stress exposure. (A) Scheme of the experiments representing the *in vitro* and *in vivo* recordings. (B) Example traces of sIPSC recordings from no odor exposed (NOE) and POSE mice. (C) Bar graphs of average sIPSC frequency (U = 11) and (D) amplitude (U = 30) in PVT/CR+ neurons recorded ex vivo from NOE (n = 8 neurons from n = 2 mice) and POSE mice 2 hours after POSE (n = 9 neurons from n = 2 mice). (E) Waveforms of three optotagged PVT/CR+ neurons recorded by the same tetrode (TT1). The neurons were recorded for two consecutive days (top vs bottom row). Channel numbers (Ch) and similarity scores (see Methods) between the two days are shown. (F-H) Alteration in the firing rate of PVT/CR+ neurons recorded for one hour both on the PRE5 (black dots) and the POST 0 days after the POSE (red dots) in the (F) wake, (G) nest, and (H) sleep states (F, t[19] = 1.499; G, t[19] = 2.871; H, t[19] = 3.378; n = 20 units from n = 4 mice). The firing rate is normalized to the mean wake population average of the PRE 5 day. (I-K) Same as E-G, for High-Frequency Activity (HFA; I, t[19] = 2.463; J, t[19] = 1.847; K, t[19] = 2.411). See Table S1 for the full results of the statistical tests. Data are shown as mean ± SEM. *p < 0.05, **p < 0.01.

To test whether this decreased inhibition can have a consequence on the post-POSE firing activity of PVT/CR+ cells we optogenetically tagged PVT/CR+ neurons and recorded their activity before POSE (PRE 5 days) and the same neuorns on POST0 day immediately after POSE (n=20 neurons from 4 animals, Fig. 3E and fig. S3, A, B). For these experiments, CR-Cre animals were injected with AAV-DIO-ChR2-EYFP (fig. S3, A) which allowed optotagging and clustering (Fig. 3E, fig. S4) of PVT/CR+ neurons using optrodes. Since many behavioral changes were confined to the nest (Fig 2 H-N) we separately analyzed firing activity during the wake state outside the nest (wake), wake state inside the nest (nest) and NREM sleep inside the nest (sleep).

We found that compared to the PRE5 day activity, on average, PVT/CR+ neurons displayed a significant increase in their firing rate during nest and sleep but not during the wake state (Fig. 3, F, G and H) on POST 0 day. Beside the mean firing rate we also analyzed the alterations in the firing pattern of PVT/CR+ neurons. Since high-frequency activity (HFA, above 100 Hz) exert disproportionally large effects on their postsynaptic targets and their synaptic plasticity^51, 52^ we quantified the occurrence of HFA (spike doublets or triplets) in the same PVT/CR+ cell population at PRE5 and POST 0 days. We found that compared to PRE5 day individual PVT/CR+ neurons displayed significantly more HFA clusters in the wake and sleep states (Fig. 3, I, J and K) immediately after the stress.

These data together show that PVT/CR+ cell activity is elevated after the stress event on the stress day, neurons display more HFA and a post-POSE reduction in inhibition can potentially be responsible for this alteration in the firing activity.

### Prolonged changes in PVT/CR+ neuronal activity after the stress exposure

In the EYFP animals behavioral alterations persisted for the entire post-stress period (POST 1-4 days, Fig 2, H-O). Thus, next, we tested, whether the observed changes in the firing activity of PVT/CR+ cells immediately after POSE (Fig 3. E-K) also persists for a more prolonged period paralleling the behavioral observations. For these experiments, we optotagged PVT/CR+ neurons (n= 69 during the PRE1-5 days and n=57 during the POST 1-4 days in 4 animals) (Fig. 4 A). Some of these neurons (n=13 tagged cells) could be recorded for up to 3 days (from PRE5 day to POST1 day (Fig. 4, D, F, H and fig. S3, B and C).

**Fig 4.**
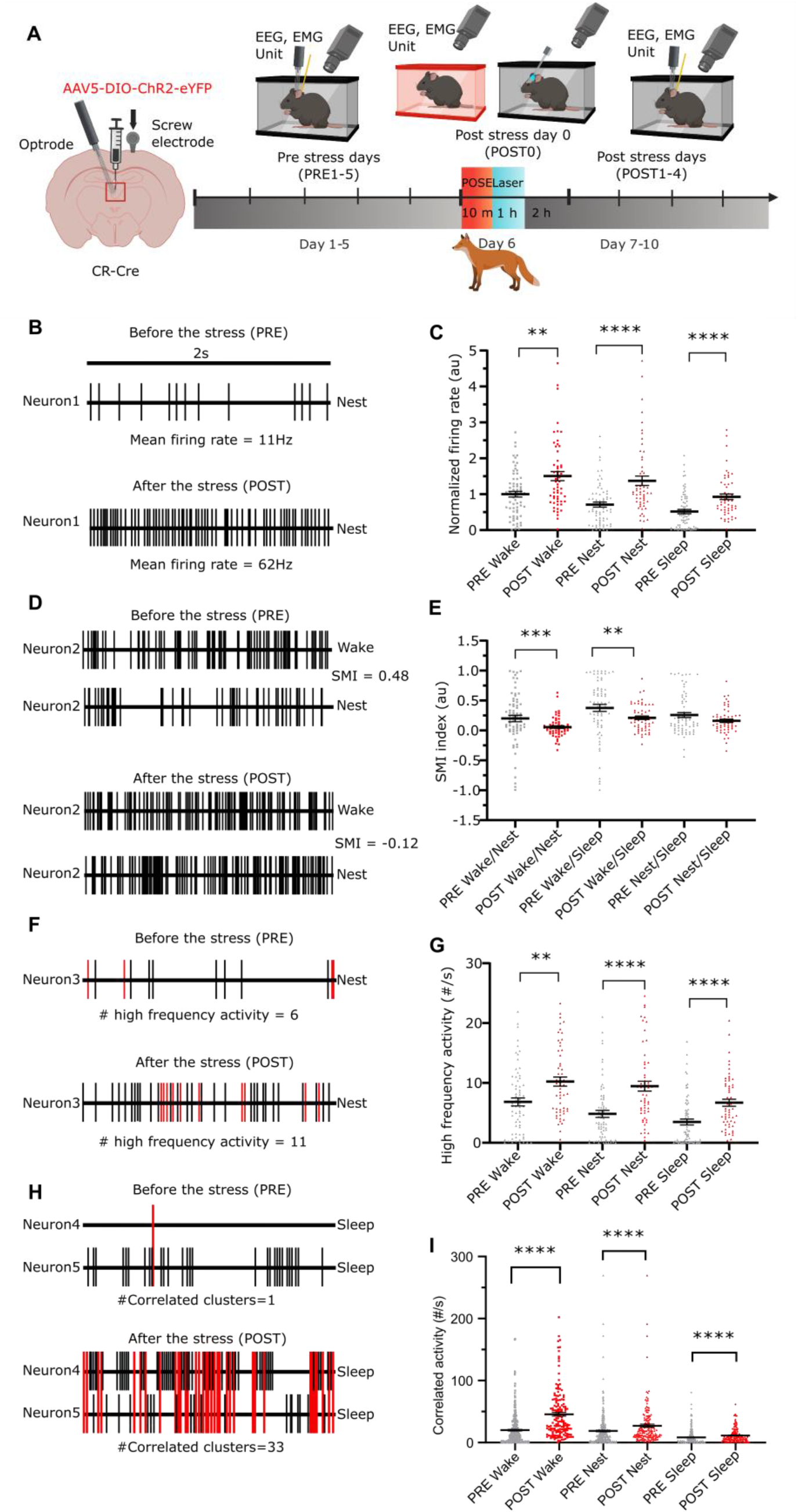
Long-term changes of PVT/CR+ unit activity after predator odor stress exposure. (A) Scheme of the experiment. (B) Change in the firing rate of an example PVT/CR+ neuron recorded on PRE 5 day and POST1 day. (C) Mean firing rate values of PVT/CR+ neurons (n = 126 neurons in 4 animals) during wake, nest, and sleep during the PRE 1-5 and the POST 1-4 days periods. Dots represent individual cells. The data are normalized to the mean wake firing rate of the PRE period (wake, U = 1357, p = 0.0026; nest, U = 1041, p < 0.0001; sleep U = 1097, p < 0.0001). (D) Alteration in the state-dependent activity of an example PVT/CR+ neuron recorded on PRE 5 and POST1 days. (E) State modulation indices (SMI) in the entire population of PVT/CR+ neurons during PRE and POST periods (wake/nest SMI, U = 1295, p = 0.0009; wake/sleep SMI, U = 1304, p = 0.0011; nest/sleep SMI, U = 1814, P = 0.4576). (F) Change in high-frequency activity (HFA) of an example PVT/ CR+ neuron recorded on PRE 5 and POST1 days. Red lines (spikes) show HFAs, black lines mark the spikes with longer interspike intervals than 10 ms. (G) Frequency of HFAs in the entire PVT/CR+ population during the entire PRE and POST periods (wake, U = 1309, p = 0.0011; nest, U = 992, p < 0.0001; sleep U = 1034, p < 0.0001). (H) Alteration in correlated activity (CA) between a representative pair of PVT/CR+ neurons recorded on PRE5 and POST1 days. Linked red lines (spikes) indicate CAs, black lines mark the spikes occurring outside the time window of 5 ms. (I) Frequency of CA in the entire population of PVT/CR+ cell pairs (n = 448 pairs) during the PRE and POST period. (U = 13625, p < 0.0001; nest, U = 20351, p < 0.0001; sleep U = 20049, p < 0.0001). See Table S1 for the full results of the statistical tests. Data are shown as mean ± SEM. **p < 0.01, ***p < 0.001, ****p < 0.0001.

During the 4 days after the stress (POST1-4), the mean firing rate of PVT/CR+ cell population significantly increased in all three states (wake, nest, and during sleep) (Fig. 4, B and C). In order to systematically compare state-dependent changes, we calculated a state modulation index (SMI, see Method) for each cell. Compared to the PRE period, SMI indices of PVT/CR+ cells significantly decreased in both the wake-to-nest and wake-to-sleep relations indicating significantly less modulation of firing rate by the behavioral state during the POST period (Fig. 4, D and E). Less state modulation was largely due to the large increase in firing rates in the resting states (nest and sleep) during the POST period (Fig. 4, C).

Next, we analyzed the alteration in firing patterns and found that relative to the PRE period the occurrence of HFA significantly increased in all three states during the POST1-4 days (Fig. 4, F, G, H, I and fig. S5, A, B). Since correlated activity (CA, spikes of two simultaneously recorded neurons within 5 ms) are also known to have a strong impact on the postsynaptic neurons we also compared the alteration in the frequency of CA between the PRE and POST periods. Similar to HFA we found that the occurrence of CA also significantly increased in all three days. After appropriate normalizations of HFA and CA with the background firing activity (see Methods) these changes remained significant only in the nest of CA values (fig. S5, C and D) indicating that an increase in the mean firing rate of PVT/CR+ neurons is largely responsible for the elevated occurrence of HFA and CA. These data together show that the activity PVT/CR+ cells display state-dependent changes persisting for days, after a single stress exposure paralleling the observed behavioral changes. Prolonged elevation of the mean firing rate of PVT/CR+ cells is mainly responsible for these alterations.

**Fig. 5.**
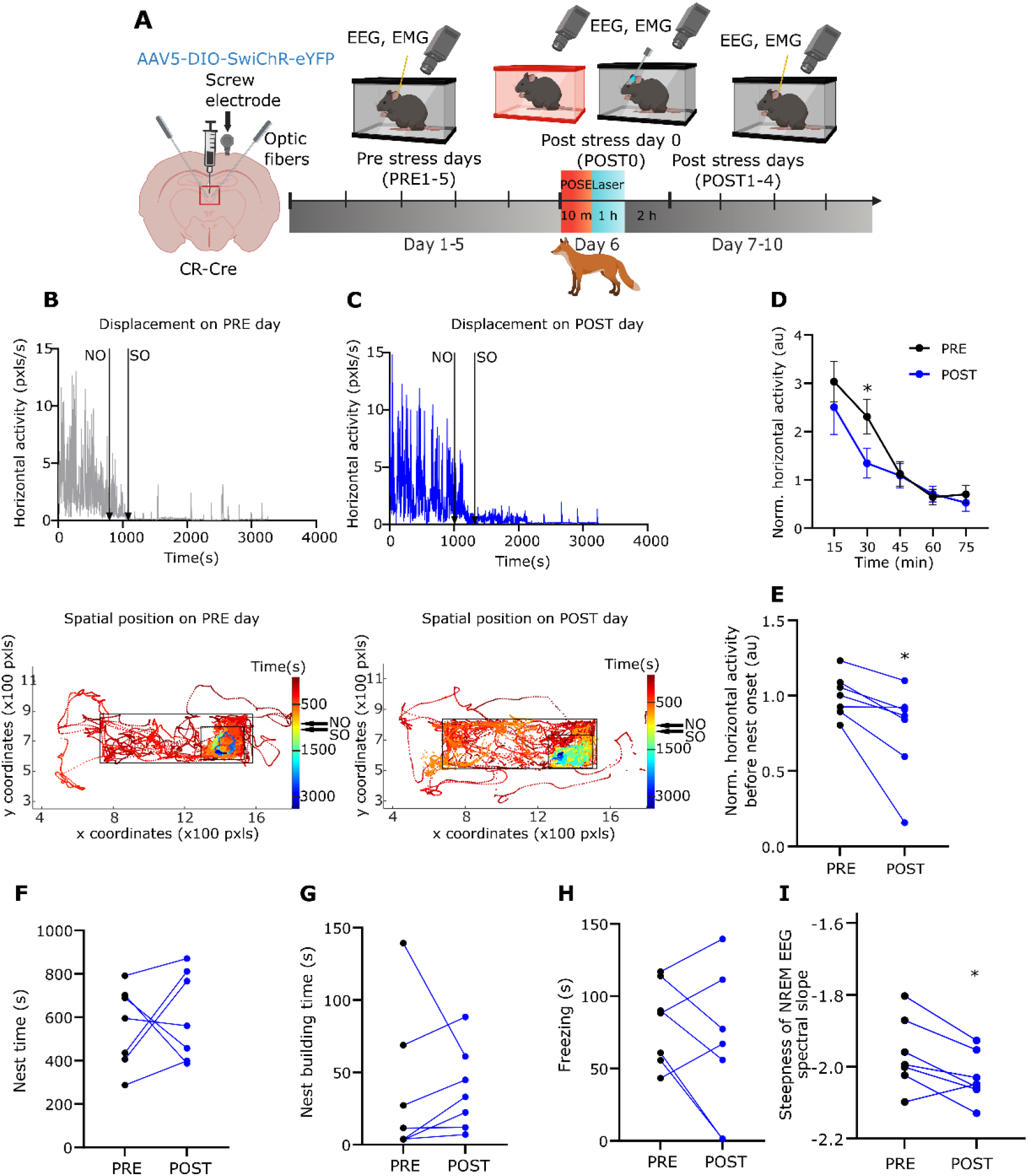
Post stress photoinhibiton of PVT/CR+ neurons prevents stress induced, long-term behavioral changes. (A) Scheme of the experiment. (B) *Top,* Changes in horizontal locomotor activity during 3 hours in one SwiChR mouse within the home cage on a PRE day. Nest onset (NO) and sleep onset (SO) are indicated with black arrows. *Bottom,* Tracked movements of the animal in the first hour. The dots show the spatial position of the head of the mouse in every frame and the color of the dots indicate lasting time (s), from red to blue; small black rectangle marks the nest area; big black rectangle marks cage area. Nest onset (NO) and sleep onset (SO) are indicated with black arrows. (C) Same as (B) on a POST day. (D) The dynamics of averaged, normalized horizontal locomotor activity (E) of SwiChR mice (n = 7) during the PRE (black) vs the POST (blue) periods (t[6] = 2.835). (E) Averaged, normalized horizontal locomotor activity of SwiChR mice during the PRE (black) vs the POST (blue) periods (t[6] = 2.944). (F) Nest time (t[6] = 0.4886), (G) nest building time (t[6] = 0.1084), (H) freezing (t[6] = 1.142) and (I) steepness of NREM EEG spectral slope (t[6] = 2.969) in SwiChR mice during the PRE and POST periods. Dots represent the averaged daily values of individual animals. See Table S1 for the full results of the statistical tests. Data are shown as mean ± SEM. *p < 0.05.

### Effect of post-stress photoinhibition on long-term changes in behavior

Since prolonged alteration of PVT/CR+ neuronal activity correlates with stress induced behavioral alterations next, we investigated whether post-stress inhibition of PVT/CR+ cells for 1 hour can prevent long-term changes in the behavior during the POST1-4 days (Fig. 5A and fig. S3, A). In contrast to EYFP animals, SwiChR mice exhibited no stress-induced behavioral changes in the investigated parameters during the POST1-4 days period compared to the PRE period. SwiChR activation resulted in decreased, not increased in locomotor activity (Fig. 5, B, C, D and E) in the POST period. Relative to the PRE period there was no alteration in the nest time (Fig. 5F) or in nest building (Fig. 5G). POSE-induced increase in freezing behavior observed in the EYFP group was also abolished in SwiChR animals for the entire POST period (Fig. 5H). Post stress inhibition significantly increased the NREM EEG spectral slope during the POST period (Fig. 5I). These data indicate that post-stress photoinhibition immediately after the stress event has prolonged beneficial effects on stress induced maladaptive effects on behavior.

### Effect of photoinhibition on long term changes in PVT/CR+ neuronal activity

Next, we investigated whether post-stress inhibition of PVT/CR+ cells for 1 hour can prevent the long-term alterations in the firing activity of PVT/CR+ cells (Fig. 6A, B) for the entire POST period similar to the behavioral changes. For these experiments we used SwiChR injected animals implanted with tetrodes. In contrast to the EYFP animals the mean average firing rate of PVT/CR+ neurons in the SwiChR animals calculated for the entire POST period (POST1-4 days) did not significantly change (n=39 neuron in 2 mice) in any state relative to the PRE period (PRE1-5 days, n=38 in 2 mice) (Fig. 6C). The marked post-POSE alteration in the state-dependent firing was also abolished. The state-modulation of neuronal activity did not decrease at the population level calculated for the entire post-POSE period (Fig. 6D). In fact, in the case of wake/sleep SMIs PVT/CR+ neurons increased not decreased their state modulation after stress in the SwiChR group (Fig 6D). Post-POSE HFA activity (Fig. 6E, fig. S6, A) and correlated activity (CA) of PVT/CR+ cells also remained unaltered during the nest and sleep state. We found only one measure, CA during the wake state, which increased significantly compared to the PRE period similar to the EYFP group (Fig. 6F, fig. S6, B).

**Fig 6.**
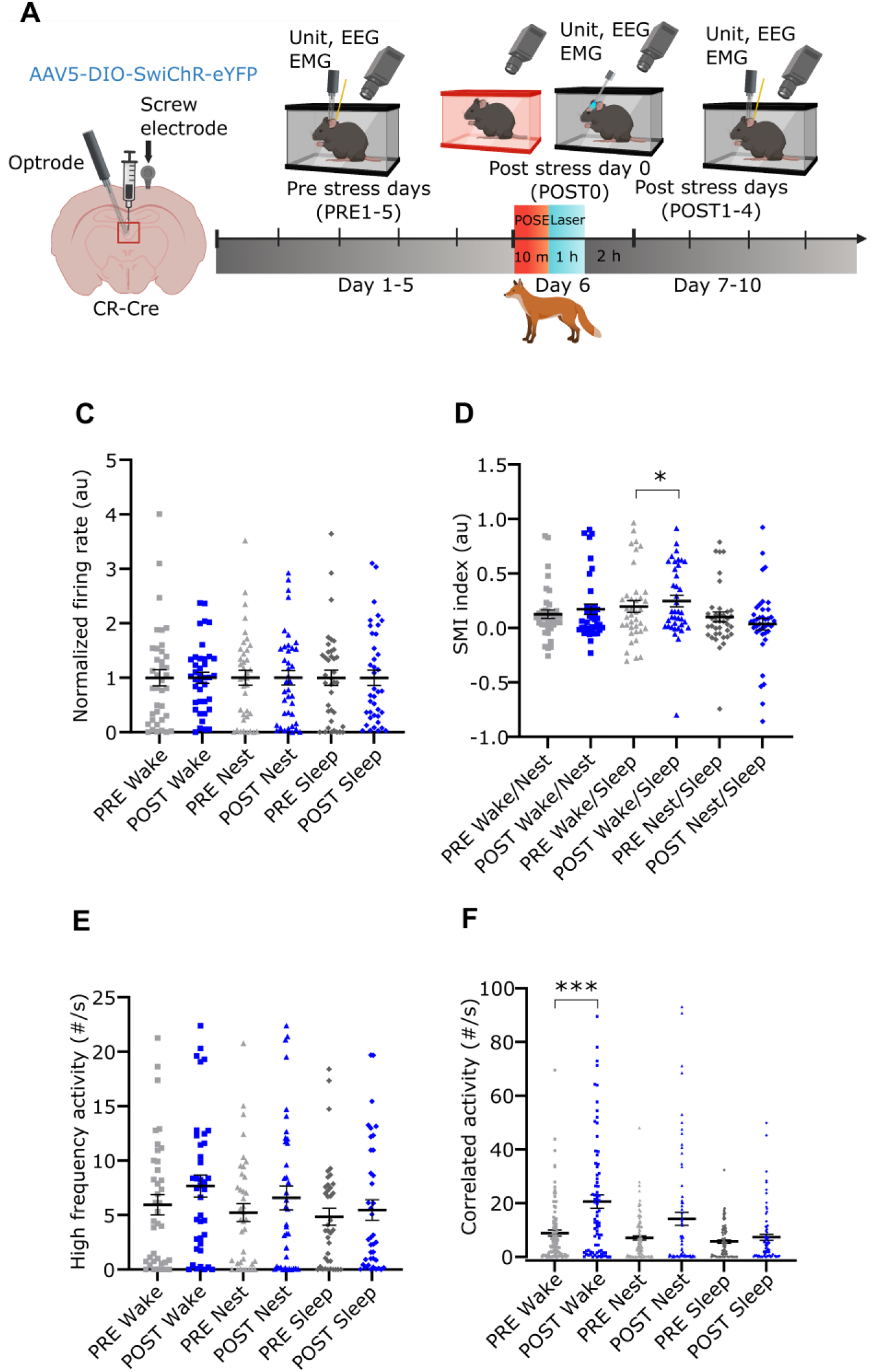
Effect of post-stress photoinhibition on long-term changes in PVT/CR+ neuronal activity. (A) Scheme of the experiment. (B) Z-scored firing rate of 9 PVT/CR+ neurons responding to SwiChR activation during the 60 min long photoinhibition protocol (see Methods). Mean (blue line) +/-SD (grey) are shown. Straight blue line above marks the laser ON time (2s). (C) Post-POSE photoinhibition of PVT/CR+ neurons on POST0 day prevents alterations in the firing rate, (D) state modulation index, (E) occurrence of high frequency activity (HFA) and (F) correlated activity (CA) during the PRE (black, n=38 neurons in 2 mice) vs the POST (blue, n=39 in 2 mice) periods (all p > 0.05 unless indicated otherwise). See Table S1 for the full results of the statistical tests. Data are shown as mean ± SEM. *p < 0.05, ***p < 0.001.

These data show that inhibiting the activity of PVT/CR+ cells immediately after stress was able to prevent not only the emergence of ASD-like phenotype but the alteration of the firing activity in PVT/CR+ neurons for the entire POST period as well.

### Role of PVT/CR+ neurons in stress induced changes of their forebrain targets

Next, we aimed to identify the impact of altered post-stress firing of PVT/CR+ neurons on their forebrain targets one hour after the POSE by measuring changes c-Fos expression in the EYFP (n=7 animals) and SwiChR (n=6 animals) groups (Fig. 7A and fig. S7). We also included the home cage (n=7) and the no odor (n=5) controls in these experiments. We selected three brain regions in which significant proportions of the subcortical glutamatergic inputs arise from PVT/CR+ cells (prelimbic cortex (PrL), basolateral amygdala (BLA), and shell of the nucleus accumbens (NAcS) (Fig. 7B) and a fourth one in which, in addition to PVT/CR+ cells, substantial glutamatergic inputs arrive from other subcortical centers (central amygdala - CeA) as well^53^. We quantified c-Fos positive neuron numbers only in regions that contained labeled PVT axons (Fig. 7B). We found that compared to homecage and NOE controls POSE markedly increased c-Fos expression in all investigated downstream brain regions (Fig. 7, C, D, E and F). Post-POSE photoinhibition of PVT/CR+ neurons was able to significantly reduce the c-Fos expression in PrL, BLA, and NAcS but not in CeA consistent with the connectivity pattern (Fig. 7, C, D, E and F). These data illustrate that altered PVT/CR+ activity immediately after the stress can lead to increased c-Fos activity in their postsynaptic targets and reducing the activity of PVT/CR+ neurons is able to alleviate the stress-induced molecular changes in key forebrain regions which control adaptive behaviors.

**Fig 7.**
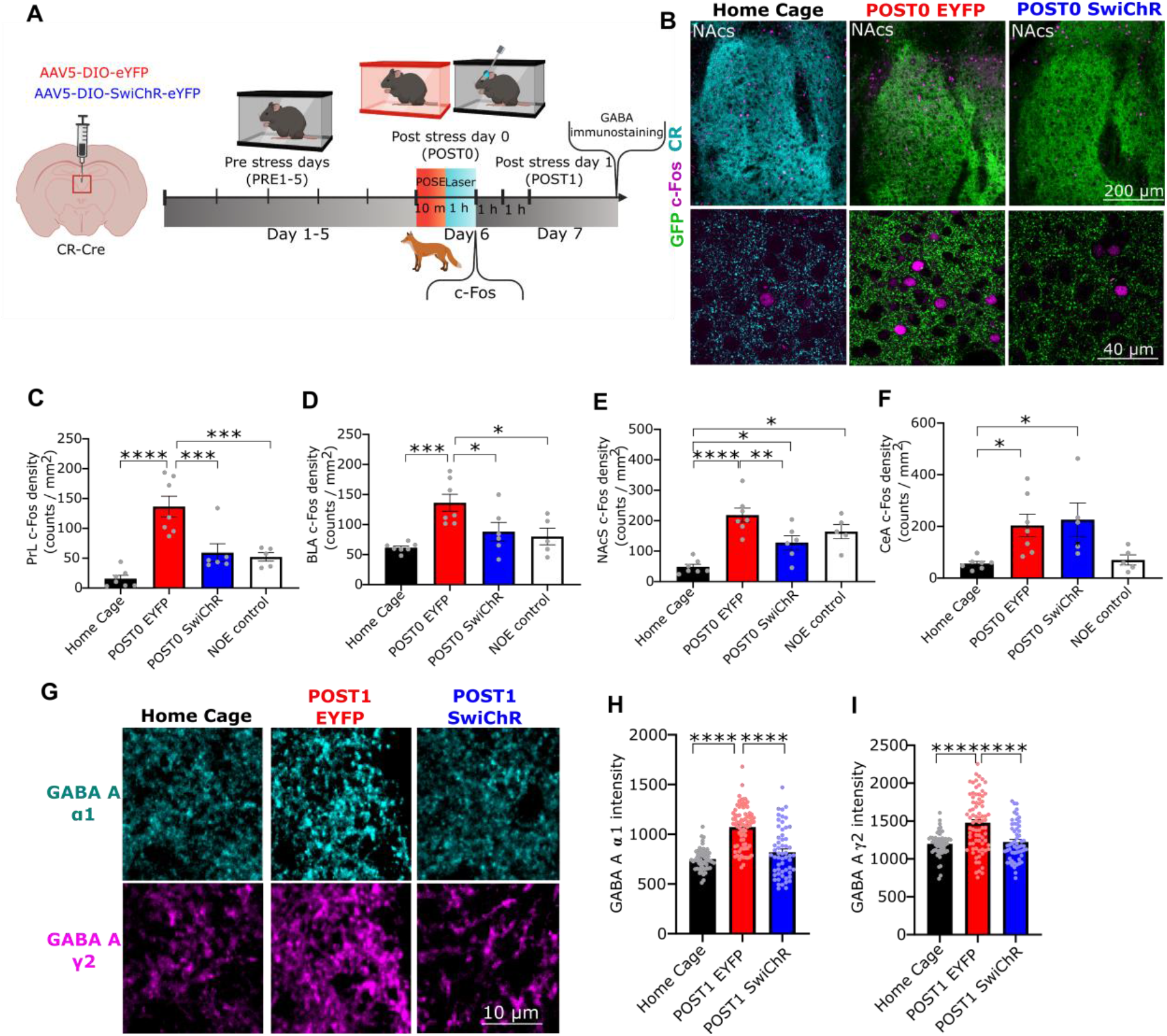
Role of PVT/CR+ neurons in stress-induced molecular changes. (A) Scheme of the experiments representing the timeline for GABA-A receptor and c-Fos immunostainings. (B) *top*) Representative, low power (upper panel) and high power (bottom panel) confocal images showing c-Fos expression (magenta) in the nucleus accumbens shell (NAcS) from Home Cage control, EYFP and SwiChR mice. Cyan (left) or green (middle and right) indicates CR-or GFP immunostaining, respectively, labeling PVT axons in NAcS. (C-F) Quantification of c-Fos positive neurons in (C) PrL, (D) BLA, (E) NAcS, and (F) CeA in Home Cage (n = 7), EYFP (n = 7), SwiChR (n = 6) and NOE (n = 5) mice on POST0 day (PrL, F(3,21) = 17.10, p < 0.0001; BLA, F(3,21) = 7.503, p = 0.0013, NAcS, F(3,21) = 14.49, p < 0.0001; CeA, F(3,20) = 4.608, p = 0.0073) (G) Representative confocal images showing GABA-A α1 and GABA-A γ2 immunostainings in the PVT from Home cage control, EYFP, and SwiChR mice on POST1 day. (H-I) Changes in the fluorescence intensity of (H) GABA-A α1 and (I) GABA-A γ2 immunostainings on POST1 day (GABA-A α1, F(2,191) = 53.04, p < 0.0001;, GABA-A γ2, F(2,191) = 21.25, p < 0.0001; Home Cage n = 59 ROI from n = 3 mice; EYFP, n = 78 ROI from n = 3 mice; SwiChR, n = 57 ROI from n = 4 mice). See Table S1 for the full results of the statistical tests. Data are shown as mean ± SEM *p < 0.05, **p < 0.01, ***p < 0.001, ****p < 0.0001.

### Stress induced changes in homeostatic GABA-A receptor expression

We observed a prolonged increase in the firing activity of PVT/CR+ neurons (Figure 5) and it is known that increased excitatory activity can lead to a compensatory homeostatic increase in synaptic inhibition^54^. Thus, using high-resolution immunocytochemistry we tested how the expression levels of the two major subunits of synaptic GABA-A receptor proteins (α1 and γ2) are altered in PVT/CR+ neurons as a result of the elevated post-POSE activity on POST1(Fig. 7A, fig. S10) in EYFP (n=3) SwiChR (n=4) and home cage control (n=3) groups. The expression of both GABA-A receptor subunits significantly increased in the EYFP animals after POSE compared to homecage controls (Fig. 7, G, H and I), demonstrating a homeostatic upregulation of GABA-A receptors due to the increased firing of PVT/CR+ neurons 1 day after the stress event. Similar to other post-stress changes post-POSE SwiChR activation was able to prevent these alterations in α1 and γ2 expression (Fig. 7, G, H and I) probably via normalizing the prolonged POST POSE increase in PVT/CR+ firing activity.

### Effects of late inhibition of PVT/CR+ cells on ASD-like phenotype

The data above strongly suggest that PVT/CR+ neurons have a critical role in establishing the acute stressed phenotype. It is still unclear, however, whether they are actively involved in maintaining the ASD-like phenotype or whether other brain regions take over this role. To gain insight into this question we applied photoinhibition not immediately after POSE but 5 days later (LATE group, Fig. 8A, fig. S8), when we still had manifest ASD-like phenotype and altered PVT/CR+ firing activity.

**Fig 8.**
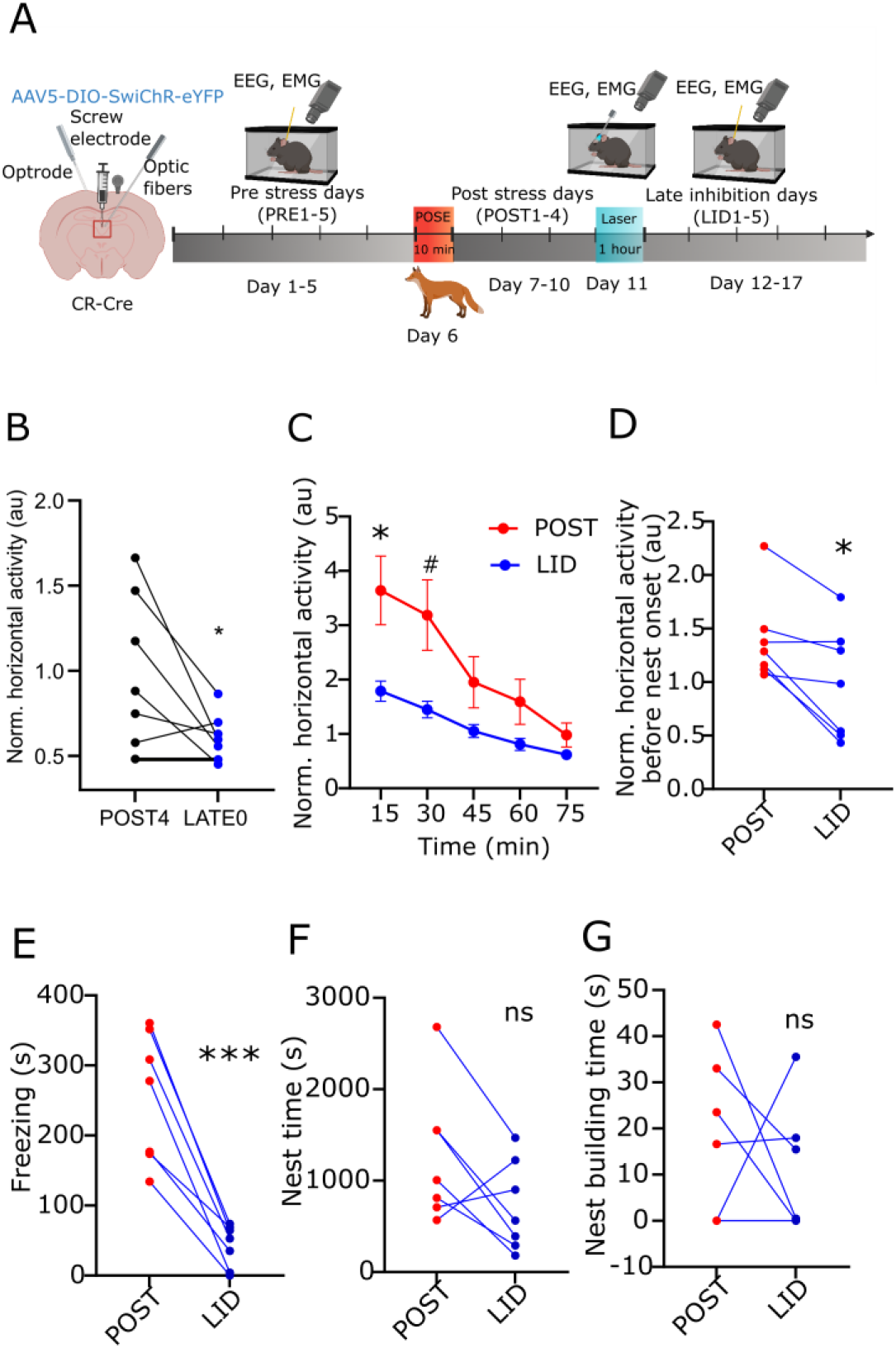
Effect of late inhibition of PVT/CR+ neurons on stress induced behavioral changes. (A) Scheme of the experiment. (B) Averaged, normalized horizontal activity during the photoinhibition in LATE0 day compared to the preceding day (POST4) (t[6] = 2.474, (n=7). (C) Dynamics of averaged, normalized horizontal locomotor activity of SwiChR mice during the POST (red) vs the LATE (blue) periods (15 min, t[6] = 2.738, p = 0.0338; 30 min t[6] = 2.443, p = 0.0503). (D) Averaged, normalized horizontal locomotor activity of SwiChR mice during the POST (red) vs the LATE (blue) periods (W = -26). (E) Freezing behavior (t[6] = 6.909, p = 0.0005), (F) nest time (t[6] = 2.031), and (G) nest building time (W = -5) in SwiChR mice during the POST (red dots) and LATE (blue dots) periods. Dots represent the average daily values of individual animals. See Table S1 for the full results of the statistical tests. Data are shown as mean ± SEM. ns means no significant difference between groups, #p < 0.06, *p < 0.05, ***p < 0.001.

Similar to P0 photoinhibition, late inhibition had an immediate effect on locomotor activity (Fig. 8, B). Late inhibition was also able to induce prolonged effects for the entire LATE period (days 12-17). Compared to the POST1-4 days horizontal activity (Fig. 8C and D) and freezing behavior (Fig. 8E) were significantly reduced in the LATE group after photoinhibition. Nest time (Fig. 8F) and nest building time (Fig. 8G), however, were not significantly altered. These data demonstrate that PVT/CR+ cells are actively involved in maintaining the ASD-like phenotype, but a single inhibitory intervention is unable to ameliorate all symptoms.

## DISCUSSION

The present data demonstrate that the post-stress firing activity of PVT/CR+ neurons immediately after the stress event is necessary for the emergence of ASD-like phenotype in mice. This conclusion is supported both by correlational and causal evidence. Neuronal activity of PVT/CR+ cells increased after the stress event in parallel with the emergence of maladaptive behavior in the entire post-stress period (POST 1-4 days) and optogenetic inhibition of PVT/CR+ neurons after the stress was able to prevent the establishment of ASD-like symptoms. Our optogenetic approach allowed us to selectively inhibit PVT/CR+ neuronal activity specifically after, but not during, the stress exposure for a well-defined time period (1 hour). In this manner we did not interfere with PVT/CR+ activity during the exposure to the stressor, thus we did not modify the perception of the stress event. Inhibition of PVT/CR+ cells after the stress event exerted long-lasting effects on both behavior and neuronal activity. A single session of one hour-long optogenetic inhibition of PVT/CR+ cells after the stress event was sufficient to prevent both the emergence of the stressed phenotype and the associated alteration in PVT/CR+ firing activity for several days. This suggests that the activity of PVT/CR+ cells in the early post-stress period is critical for the establishment of behavioral alterations induced by stress.

In this study, we proposed and validated a rodent stress model for human ASD. The stressor was an inescapable threat (predator odor) based on innate fear. In agreement with the literature^39–41, 43^ strong stress response to 2MT was demonstrated by robust defensive behavior during and immediately after the exposure, elevated c-Fos expression in PVN, and a significant increase in blood corticosterone levels after the POE. In order to capture ASD-like alterations in the peritraumatic time window (5-5 days) we used multiday recordings of behavior and neuronal activity and systematically compared pre vs. post-stress data. Human ASD involves a complex alteration of behavior which consists of several entirely human-specific symptoms^2^ but some have their rodent equivalents. Our demonstrated significant alteration in locomotion, freezing in the nest, nest-building behavior, and the sleep onset for several days after the stress that prove the etiological and face validity of this animal model by mimicking well some of the core symptoms of human ASD like hypervigilance, intense emotional distress, and sleep disturbance.

The involvement of PVT/CR+ neurons in stress induced alteration of behavior is consistent with earlier behavioral data^31^ as well as with the known connectivity of PVT/CR+ neurons ^6, 7, 20^. These earlier as well as our present data suggest that PVT/CR+ neurons represent a critical bottleneck between the brainstem and forebrain centers concerning the transmission and distribution of stress-related signals to the forebrain^17^ . The similarity in the organization of PVT between mice and humans^7^ and the involvement of human PVT in arousal-related problems^27^ suggest that PVT may play a similar role in humans.

Our data highlight the sensitivity of pre-sleep behavior in the nest to stress. Before the stress exposure, the animals displayed normal nest building, spent little time freezing in the nest, and fell asleep relatively fast after they entered the nest. During these prestress days, the firing rate of PVT/CR+ cells in the nest (while the animals were still awake) was significantly slower than outside the nest but faster than during sleep. These behavioral and physioloigcal data support the view that the wake state in the nest is a transient arousal state and represents the preparation to sleep. Following the stress exposure, both the behavior and the firing activity in the nest changed significantly. Mice displayed more freezing but less nest-building behavior and needed more time to fall asleep. In parallel, the firing activity of PVT/CR+ cells in the nest was also significantly altered. In contrast to pre-stress days, during the post-stress days the average wake/nest SMI values of PVT/CR+ cells were close to zero (0.05, Fig 4E) indicating that, as a population, the neurons did not any longer slow down upon entering the nest on the post-stress days. These data indicate that similar to humans, pre-sleep behavior is a highly sensitive to stress and the post-stress elevation of PVT/CR+ neuronal activity in the nest contributes to the altered nest behavior.

We observed increased c-Fos activity in the forebrain targets of PVT/CR+ neurons (PrL, Amy, nAC) after the stress exposure. This is consistent with the facts that PVT/CR+ cells are able to evoke strong postsynaptic response in these brain areas^7^ and that PVT/CR+ neurons increased their activity after the stress exposure (present study). The high-frequency activity (<100 Hz) of PVT/CR+ was also significantly elevated after the stress. Since these events have a large impact on the postsynaptic targets^52^, they can have the capacity to induce altered activity in the forebrain. Inhibiting PVT/CR+ neurons after POSE prevented the increase of c-Fos activity in the postsynaptic targets within the terminal field of the SwiChR labelled PVT/CR+ cells. This confirms the critical role of PVT/CR+ cells in the stress induced increase in c-Fos activity in PrL, Amy and nAC. These data together suggests that elevated PVT/CR+ neuronal activity after the stress event can lead to altered behavior via acting on key forebrain centers^5, 45, 55^. Our data shows that increased PVT/CR+ firing rate may be the consequence of reduced synaptic inhibition immediately after the exposure to stress. This is consistent with earlier data, showing a D2 receptor-mediated decrease in inhibition after the stress^31^, It has to mentioned though that in our case the amplitude of spontaneous IPSCs was not reduced, which may be explained by the different stressor used.

In this study, the elevation of the firing rate was not confined to the period immediately after the stress. Rather, PVT/CR+ neurons maintained altered firing rates and patterns for several days after the stress event. This suggests that the sustained increase of PVT/CR+ firing activity may be the mechanism that supports the establishment of ASD. Sustained increase in firing activity is rarely observed in the brain. The firing rates of individual neurons are maintained by a cell autonomous form of homeostatic plasticity (called “synaptic scaling”)^56–58^. Shift in neuronal activity has been demonstrated before ^59, 60^ but in these cases, the perturbed firing rates of individual neurons gradually returned to a precise, cell-autonomous set point^61^. In our case, however, elevated activity of PVT/CR+ cells persisted for days. Increased neuronal activity can also lead to a homeostatic upregulation of GABA-A receptor density^54^. We also found that in parallel to the elevated unit activity expression of synaptic GABA-A receptors subunits was significantly increased in PVT/CR+ cells after the stress. Apparently, however, in our case, increased synaptic inhibition was not sufficient to reduce the elevated post-stress firing rate of PVT/CR+ cells. Morever, since GABAergic mechanisms are known to be involved in the synchronization of neuronal activity^62, 63^ this homeostatic increase in inhibition can actually be responsible for the prolonged increase in synchrony observed in the post-stress period among PVT/CR+ cells. Mechanisms of persistent activity in the fear circuit are intensively studied^64^. Understanding the cellular bases of the post-stress shift in PVT/CR+ activity requires further investigation.

Optogenetic inhibition of PVT/CR+ cells resulted in a clear reduction of firing rate during the laser ON periods. This SwiChR activation reduced the stress-induced increase in c-Fos activity in PVT/CR+ cells confirming the inhibitory effect of the opsin, especially around the optic fibers. Diminishing the activity of PVT/CR+ neurons for 1 hour after the stress event had a prolonged effect on both PVT/CR+ activity and behavior. The optogenetic interference with post-stress neuronal activity was able to block both the shift in the firing regime of PVT/CR+ induced by stress for the whole duration of observations (5 days) and the emergence of ASD-like symptoms. In agreement with these, post-stress upregulation of GABA-A receptors also did not take place in the SwiChR group. These data indicate that the optogenetically evoked Cl^-^ entry via the SwiChR molecule during the early post-stress period is able to counteract the mechanisms which finally lead to the prolonged stress-induced shift in PVT/CR+ excitability.

Inhibition of PVT/CR+ neurons five days after stress was still able to alleviate some (but not all) symptoms of ASD suggesting that after five days the maintenance of the whole symptom cluster is not entirely dependent on the persistently elevated PVT/CR+ activity any longer. Rather, by this time apparently some symptoms have already consolidated in the forebrain targets as a consequence of the prolonged excitatory bombardment of PVT/CR+ cells.

## Acknowledgement

We thank the Light Microscopy Center and the Virus Technology Unit at the Institute of Experimental Medicine for kindly providing microscopy support. Authors would like to thank Krisztina Faddi, Dániel Kuti, Kornél Demeter and Csaba Dávid for their excellent technical assistance. The schemes were created with www.biorender.com.

## Funding

ERC Advanced Grant FRONTHAL, 742595 (LA)

ERC Starting Grant CholAminCo, 715043, (BH)

National Research, Development and Innovation Office NKFIH K135561, (BH)

European Union project within the framework of the Artificial Intelligence National Laboratory RRF-2.3.1-21-2022-00004 (LA)

The Ministry of Innovation and Technology of Hungary within the framework of the Higher Education Institutional Excellence Program (TKP2021-EGA-25) (RB)

ANR-21-CE37-0025-02, ANR-21-CE16-0012-02 (MAD)

## Author contributions

Conceptualization: AJ, LB, LA

Methodology: AJ, LB, LA, BK, BH, OSZ, RB

Investigation: AJ, LB, ZSB, HK, KK, GK, MD

Visualization: AJ, BK, BL

Funding acquisition: LA, BH, KK, RB.MD

Project administration: LA

Supervision: LA

Writing – original draft: AJ, LB, LA

Writing – review & editing: AJ, LB, BH, MD, KK, RB, GK

## Data and materials availability

Individual data points used to create the figures are available as Source Data Files. All raw data and custom codes that support the findings, tools and reagents will be shared on an unrestricted basis; requests should be directed to the corresponding authors. Concerning the data we are able to provide the following datasets upon request: images of the full extent of the injection sites, raw Open Ephys files of individual PVT/CR+ cell and cortical LFP activities, raw in vitro data and video recordings.

## Supplementary Materials

Materials and Methods

Figs. S1 to S12

Movies S1 to S6

Tables S1

## STAR METHODS

All procedures with mice were approved by the Animal Welfare Committee of the Institute of Experimental Medicine, Budapest, conformed to guidelines established by the European Community’s Council Directive of November 24, 1986 (86/609/EEC). Experiments were authorized by the National Animal Research Authorities of Hungary (PE/EA/877-7/2020).

### Subjects

Adult male *Calb2-IRES-Cre* (CR-Cre, C57Bl/6J, n=73) mice (postnatal 3-5 months) were used in the experiments from the Jackson Laboratory (RRID: IMSR_JAX:10774). All animals were group-housed (three to four / cage) in Plexiglas chambers at constant temperature (22±1°C) and humidity (40–60%) under a circadian light-dark cycle (lights-on at 9 A.M., lights-off at 9 P.M.). All mice were single housed after electrode and optic fiber implantation. For the animals the food (Sniff) and water were provided *ad libitum*.

### Viral vectors

For control mice in the behavioral, sleep, corticosterone, c-Fos and GABA-receptor localization experiments we used AAV5.EF1a.DIO.eYFP.WPRE.hGH (Addgene27056) vector (EYFP mice), whereas for optogenetic inhibition in the same experiments, AAV5.EF1a.DIO.SwiChRCA.TS.EYFP.WPRE (UNC Vector Core) vector was injected (SwiChR mice). In the optrode experiments for the not inhibited, control animals (n=4 ChR2 animals) we utilized AAV5.EF1a.DIO.hChR2(H134R)-eYFP.WPRE.hGH (Addgene20298P) vector for optical tagging, whereas for the inhibited animals with unit recordings the same SwiChR construct was utilized to phototagging as above (n=2 SwiChR animals).

### Surgery

Mice were deeply anesthetized with ketamine-xylazin (intraperitoneal injection, ketamine, 83 mg/kg; xylazine, 3.3 mg/kg). Depth of sleep was monitored throughout the surgery, if needed supplemental dose (ketamine, 28 mg/kg; xylazine, 1.1 mg/kg) was injected intramuscularly. Externally applied lidocaine was used on the head and ears. The mice were placed in a stereotaxic frame (Kopf Instruments, Tujunga, CA) on a heating pad to prevent hypothermia. We used eye protection gel against drying. The scalp in the midline was removed (from Bregma to Lamda) and the skull thoroughly cleaned and treated with 3% hydrogen peroxide for implantation.

#### Viral injection

A craniotomy was drilled above the PVT and the virus was slowly injected (1nl/min) into the PVT (AP: -0.9, ML: 0, AP and ML taken from the bregma). We injected 50-50 nl in 2 dorsoventral positions (DV: 3.1 and 2.9, DV taken from the brain surface) in the behavioral, corticosterone, c-Fos and GABA-A receptor localization experiments and 150-150 nl in the same positions in case of the tetrode experiments. Virus was allowed to express for a minimum of 3 weeks before behavioral and tetrode experiments (fig. S2, A, fig. S3, A, fig. S7, fig. S8).

#### Implantation

In the behavioral, corticosterone, c-Fos and EEG experiments two optic fibers were implanted above the PVT (AP: -0.9 and -1.4, ML: 0, DV: 2.8, 10° angle), together with three EEG screw electrodes (2 above the frontal lobe, at AP: +0.2, ML: +/-1.8 and 1 above the parietal lobe at AP: -1.1, ML: +/-1.8) and one EMG wire electrode in the neck muscle (directly behind the skull). The EEG and EMG electrodes were soldered to Omnetics connectors (Omnetics). In the tetrode experiments tetrodes were also implanted above the PVT (AP: -0.9, ML: 0, DV: 2.7, 9.5°angle), and gradually screwed down to PVT until the start of the experiment. Optic fibers, EEG and EMG electrodes were fixed with dental cement (Metabond), tetrode wires were fixed with flexible glue first and then also with cement (Fig. 4A, Fig. 5A, Fig. 6A). After surgery, the animals were given saline and painkillers for 1-5 days according to their condition. Recovery from surgery lasted 2 weeks.

#### Recording apparatus

All mice were recorded in single-animal boxes with sound and electrical noise isolation. Cameras (FlyCapture) and LEDs were built in the boxes, providing an ambient 200 lux (50 cm from the cage) light. Electrophysiological recordings were provided via Intan Interface Board, Intan cables and chips (Intan Technologies). To avoid entanglement of wires and fibers optics as well as to reduce the weight of the cables and connectors a dual (wire and fiber optic) commutator system was used (Imetronic), connecting to the Intan based recording system and the head implants of the animals. We recorded the electrophysiological signal with a custom-made script of the Bonsai visual-programming software.

### Predatory Odor Stress Exposure (POSE)

We assessed defensive behavior to an ecologically relevant aversive stimulus, i.e. predator odor stress exposure (POSE) by using a synthetic analog of a fox anogenital product (2-methyl 2-thiazoline; 2MT), in a transparent plexiglass arena (32 x 12.5 x 15 cm). Testing was carried out in a fume hood with high intensity light (280-320 lux) in covered cages to equalize odor exposure across subjects. At start, mice placed in the testing cage and were left to freely explore it for 2 min which also served as a baseline period. Each subject received fox odor (2-MT) presentation for 10 min ^40, 41^. 2 µl pure, undiluted 2-MT (sc-251779, Santa Cruz Biotechnology) was pipetted onto a filter paper (2 x 2 cm) which was put into an Eppendorf tube (1.5 mL). After a 2 min baseline period, the Eppendorf was placed into the left corner of the test cage. Mouse behavior was recorded (Bonsai) and quantified (Solomon) using a video-based measurement system. We used two behavioral variables to characterize the innate behavioral responses: freezing and escape jumps. The mice were considered to freeze if movement was not detected for 2 s. Identical procedure was used for the no odor exposure (NOE) experiments, but in this case 2-MT was omitted. In case of home cage controls the animals were left in their home cages until the beginning of the perfusion.

#### Behavioural protocol

In the behavior experiments we distinguished three major states: the pre-stress period (5 days PRE 1-5 days), the day of POSE (POST 0 day) and the post-stress period (4 days POST 1- 4 days) (Fig. 1A, Fig. 2A and Fig 3A). In case of the LATE group, we made the optogenetic inhibition 5 days after the POSE and then we continued recording in the next 5 consecutive days, the same way as detailed above (Fig. 8A).

Animals were habituated to the recording apparatus for 3 days (3 hours/day) prior to the experiments. The home cages of the animals were placed to sound insulated boxes, their lids were removed, and the cables were connected to the head implants of the animals. Animals were allowed to move freely and to climb to the edge of their home cages. During the pre-stress periods the baseline activity of the mice (EEG /EMG and behavior in n=28 animals, PVT/CR+ unit activity and behavior in n=6 animals) were recorded for three hours every day starting at the beginning of their light phase (ZG time 0) for five days. On the day of the POSE animals were transferred to a new room and were placed into a new cage for POSE (see above). After POSE we put the animals back to their come cages and continued recording for the next 3 hours. In the first hour the laser protocol was performed, which meant inhibition in case of the SwiChR group. On the POST period we recorded the activity of the mice in the same way as on the PRE period. POST period lasted for four days.

#### Tetrode recording

In the tetrode experiments movable drives were built and implanted. 28 channels of unit activity, 1 channel of EEG and 1 channel of EMG were recorded (Fig. 4A, Fig 5A and Fig 6A).

The tetrode drives consisted of 7 tetrodes (28 channels), an optic fiber (105 µm core diameter, 0.22 NA, Thorlabs, New Jersey, US), an EMG wire electrode, EEG screw electrode, a ground and a reference cable attached to the electrode interface board. The tetrodes were positioned 300 micrometers below the tip of the optical fiber. The end of the optical fiber was milled into a cone shape. The tetrodes were gold-plated the day before implantation to a resistance below 80 k ohms

In tetrode recording animals were subjected to the same behavioural test as described above (see in behavioural protocol section). Phototagging was used to identify PVT/CR+ cells (see in optogenetic inhibition and phototagging section). Data were recorded with OpenEphys board and software. Video was recorded in the same way as in the behavioural experiments (see in behavioural protocol section).

### Optogenetic inhibition and phototagging

In all experiments, immediately after the POSE the animals were returned to their home cage and two external optic fibers (105 µm core diameter, 0.22 NA, Thorlabs, New Jersey, US) were coupled to the implanted optic fibers with a ceramic sleeve. A rotary joint (Doric Lenses, Québec, Canada) was built in the commutator system to avoid the coiling of optic fibers and Intan cables. Optical manipulations were provided with a 473 nm DPSS laser (LaserGlow Technologies). Before every use laser intensity was checked by a manual photometer (Thorlabs, New Jersey, US). Laser intensity was also monitored online during the stimuli via built in photosensors (Thorlabs, New Jersey, US). Lasers were controlled via Matlab with a National Instrument Board (Matlab) and Arduino (Arduino Uno). Photoinhibition was maintained by laser pulses and the same laser protocol was performed in case of the SwiChR group (stressed and inhibited) and also in EYFP group (stressed) for 60 minutes. Both for EYFP (stressed, n=43 mice) in SwiChR (stressed and inhibited n=30 mice,) mice trains of 2 s ON and 13 s OFF light pulses were used (10-15 mW).

For phototagging in ChR2-expressing mice (n=4) we utilized 4 Hz, 1ms long pulses (0.5- 5 mW; 120-150 stimulation) at the beginning and at the end of the recordings (fig. S4, D). For phototagging in SwiChR animals (n = 2) we used four, 2 s long (2 s ON and 13 s OFF) pulses (0.5-5mW) during wakefulness as well as during sleep (4-4 trains) (fig. S1).

### Histological verification of viral infection and optic fiber/tetrode placement

Mice were anesthetized with 2% isoflurane followed by an intraperitoneal injection of an anesthetic mixture (ketamine, 83 mg/kg; xylazine, 3.3 mg/kg) to achieve deep anesthesia. Next, mice were perfused transcardially with 0.9% saline for 2 min followed by 100 ml of 4% PFA solution. After perfusion, brains were removed from the skull, and thalamic section were immersion-fixed in 2% PFA for 30 min. Brains were cut into 50 μm sections using a vibrating microtome (Vibratome 3000). To verify the viral infection and cannula/fiber optic placement every 3^rd^ sections were stained from each animal. Sections were washed four times in 0.1 M phosphate buffer (PB) and incubated in a blocking solution containing 2% TritonX-100 and 10% normal donkey serum (Jackson ImmunoResearch, USA) in PB for 2 hours at room temperature.

Sections were then incubated (24 hours) with anti-calretinin guinea pig (1:1000, 214104, Synaptic Systems, RRID: AB_10635160), anti-GPF chicken (1:1000, A10262, Thermo Fisher Scientific, RRID: AB_2534023) and anti-IBA1 rabbit (1:500, 27030, Wako, RRID:AB_2314667) primary antibodies dissolved in 2% normal donkey serum solution. Afterward, sections were washed in PBS 4 times (10 minutes each time), then incubated with secondary antibodies. The primary antibodies were visualized using the following secondary antibodies: Alexa Fluor 488-conjugated donkey anti-chicken (1:1000, 703-545-155, Jackson ImmunoResearch), Cy3-conjugated donkey anti-rabbit (1:1000, 711-165-152, Jackson ImmunoResearch), and Alexa Fluor 647 conjugated donkey anti-guinea pig (1:1000, 706-605-148, Jackson ImmunoResearch) antibodies. After being washed with PBS (phosphate buffer saline), the sections were mounted on glass slides and cover slipped using Mowiol 4–88 (81381, Sigma-Aldrich) fluorescent mounting medium. Images were acquired on Panoramic Digital Slide Scanner (Zeiss, Plan-Apochromat 10X/NA 0.45, xy: 0.65 μm/pixel, Panoramic MIDI II; 3DHISTECH, Budapest, Hungary).

#### c-Fos mapping with corticosterone measurement

Following POSE, neuronal activation and stress hormone reactivity were investigated using c-Fos immunohistochemistry and serum corticosterone measurement, respectively. The following experimental groups were included:

*1) Home cage control group* (Home Cage, n=7). These animals were left in their home cage for the whole duration of the experiment. They were not moved to a new cage on the day of the experiment, received no POSE or laser stimulation.
*2) No Odor Exposure (NOE) group* (n=5). These animals were moved to the new cage, were not exposed to 2 MT there but received laser stimulation when they returned to their home cage using the same protocol as the other two groups.
*3) EYFP group* (n=7). These animals were moved to the new cage, were exposed to 2 MT and received laser stimulation when they returned to their home cage. Since these animals were injected with EYFP containing virus PVT/CR+ cell did not experience optogenetic inhibition.
*4) SwiChR group* (n=6). These animals were moved to the new cage, were exposed to 2 MT and received laser stimulation when they returned to their home cage. The activity of PVT/CR+ cells were reduced by optogenetic inhibition (Fig 6B).

All groups were submitted the same habituation protocol to familiarize the mice with the recording environment (for details see behavioural protocol section). Mice were decapitated 60 min after POSE. Trunk blood was collected for CORT radioimmunoassay (RIA) and brains were fixed in 4% PFA for 3 days. c-Fos, CR, GFP triple staining was performed on 50 μm thick coronal sections selected by a 1:6 fractionator sampling covering the whole rostral to caudal extension of PVT and the forebrain regions targeted by PVT/CR+ neurons including prelimbic cortex (PrL, +2.22 mm to +1.54 mm from Bregma), nucleus accumbens shell (NAcS, +1.94 mm to+ 0.86 mm from Bregma), paraventricular hypothalamic nucleus (PVH, –0.58 mm to –1.22 mm from Bregma), central amygdala (CeA, –1.22 mm to –1.94 mm from Bregma), and basolateral amygdala (BLA, –0.58 mm to –1.94 mm from Bregma).

For immunocytochemistry, free-floating brain sections were washed in 0.1 M PB and blocked in 10% normal donkey serum (NDS) in PB containing and 2% Triton-X 100 for 2 hours at room temperature. The primary antibodies against c-Fos (rabbit, 1:1000, 226003, Synaptic Systems, RRID: AB_2231974), calretinin (guinea pig, 1:1000, 214104, Synaptic Systems,

RRID: AB_10635160) and GFP (chicken, 1:1000, A10262, Thermo Fisher Scientific, RRID: AB_2534023) were diluted in PB containing 2% NDS. Twenty-four hours later, sections were washed multiple times and were incubated with Alexa Fluor 488-conjugated donkey anti-chicken (1:1000, 703-545-155, Jackson ImmunoResearch), Cy3-conjugated donkey anti-rabbit (1:1000, 711-165-152, Jackson ImmunoResearch), Alexa Fluor 647-conjugated donkey anti-guinea pig (1:1000, 706-605-148, Jackson ImmunoResearch) antibodies and with Hoechst 33258 (1:2000, Sigma-Aldrich) for 2 h at room temperature. After further PB washes, sections were mounted on glass slides and coverslipped using Mowiol 4–88 (81381, Sigma-Aldrich) fluorescent mounting medium. Images were taken using a Panoramic Digital Slide Scanner (Zeiss, Plan-Apochromat 10X/NA 0.45, xy: 0.65 μm/pixel, Panoramic MIDI II; 3DHISTECH, Budapest, Hungary). Every 3rd section was stained and analyzed per animals. To assess c-Fos activation in the PVT/CR+ neuron projections targets, we delineated and identified regions of interest (PrL, NAcS, BLA, CeA) based axonal labelling and on neuroanatomical landmarks on fluorescent pictures using CaseViewer 2.3 software (3DHISTECH). c-Fos signal was counted bilaterally in covering the whole antero-posterior extension of the actual region/subregion.

To assess the POSE induced activation of PVT/CR+ neurons, c-Fos immunoreactivity of CR expressing cells were analyzed manually in 20 × confocal images below the optic fiber using a Nikon A1R confocal microscope (CFI Plan Apo VC20X/N.A. 0.75, xy: 0.31 μm/pixel, Nikon Europe, Amsterdam, The Netherlands).

To quantify c-Fos density throughout the rostrocaudal extension of the PVT, anti-c-Fos (rabbit; 1:20000, PC38, Calbiochem, RRID: AB_2106755) was developed with DABNi as a chromogen. All sections used for quantification were developed together for the same duration. The section was dehydrated and then mounted with DePex (Serva, Heidelberg, Germany). The number of c-Fos-labeled cells was analyzed using a custom-written ImageJ (RRID:SCR_003070) script.

#### Corticosterone measurement

Serum corticosterone was measured by direct RIA as described ^65^. Briefly, the corticosterone antibody was raised in rabbits against corticosterone-carboxymethyloxime bovine serum albumin conjugate. ^125^I-labeled corticosterone-carboxymethyloxime-tyrosine methyl ester was used as tracer. The interference from plasma transcortin was eliminated by inactivating transcortin at a low pH. Assay sensitivity was 1 pmol, the intraassay coefficient of variation was 7.5%.

### GABA-A receptor subunit expression analysis

To investigate the effect of stress on GABAA receptor subunit expression levels in the PVT, we used SwiChR (n=3) and EYFP (n=3) injected animals which underwent POSE as described previously (Fig. 4A and fig. S7). 24 hrs after POSE, injected and not injected homecage control (Home Cage, n= 3) mice were anesthetized with 2% isoflurane followed by an intraperitoneal injection of an anesthetic mixture (ketamine, 83 mg/kg; xylazine, 3.3 mg/kg) to achieve deep anesthesia. Next, mice were perfused transcardially with 0.9% saline for 2 min followed by 100 ml of 1% PFA solution. After perfusion, brains were removed from the skull, and thalamic section were immersion-fixed in 2% PFA for 30 min. Brains were cut into 50 μm sections using a vibrating microtome (Vibratome 3000). Perfusion-fixed sections were washed four times in 0.1 M phosphate buffer (PB) for 1 hour. For detection of GABAA-receptor γ2 and α1 subunits, sections were pretreated with 0.2 M HCl (hydrogen-cloride) solution containing 0.2 mg/ml pepsin (Dako) at 37°C for 2 min, followed by a 2 hours long blocking period in 10% normal donkey serum in PB ^66^. This was followed by an overnight incubation at room temperature in a solution containing 2% NDS, 0.05% sodium azide and a mixture of primary antibodies: rabbit anti-γ2 (1:2000, 224003, Synaptic System, RRID:AB_2263066) guinea-pig anti-α1 (1:500, GABAARa1-GP, Frontier Institute, RRID:AB_2571572) and chicken anti-GFP (1:1000, A10262, Thermo Fisher Scientific, RRID:AB_2534023) to enhance the endogenously expressed EYFP signal. The primary antibodies were visualized using the following secondary antibodies: Alexa Fluor 488-conjugated donkey anti-chicken (1:1000, 703-545-155, Jackson ImmunoResearch), Cy3-conjugated donkey anti-rabbit (1:1000, 711-165-152, Jackson ImmunoResearch), and Alexa Fluor 647 conjugated donkey anti-guinea pig (1:1000, 706-605-148, Jackson ImmunoResearch) antibodies. After being washed with PBS (phosphate buffer saline), the sections were mounted on glass slides and cover slipped using Mowiol 4–88 (81381, Sigma-Aldrich) fluorescent mounting medium. Images from PVT were obtained using a Nikon A1R confocal microscope (Plan Apo VC 60X oil objective; numerical aperture, 1.40; xy, 0.10 µm/pixel; z-step size, 0.10 µm). Images were taken through a z-plane (1.5 µm) at the center of the tissue containing 20 stacks within each region. During image acquisition gain, offset, laser intensity, zoom, and pinhole were kept constant. In each animal, 12–30 regions of interest (ROIs, rectangular frame) restricted to the PVT were analyzed from 3 brain sections. To determine the expression levels of GABAA-receptor subunits, the images were thresholded and gray values of pixels were registered in ROIs defined in appropriate optical sections of image stacks in ImageJ software (RRID:SCR_003070).

### Ex vivo slice preparations

Ex vivo electrophysiology was performed on adult CR/IRES-Cre mice (n = 4) previously infused in the PVT with the AAV5.EF1a.DIO.SwiChRCA.TS.EYFP.WPRE virus. Coronal brain slices (300 μm) including the PVT were prepared minimum 4 weeks following surgery. Following anesthesia with isoflurane, the brain was quickly removed in an ice-cold bicarbonate buffered (BBS) solution containing the following (mM): 116 NaCl, 2.5 KCl, 1.25 NaH_2_PO_4_, 26 NaHCO_3_, 30 glucose, 1.6 CaCl_2_, 1.5MgCl_2_, and 5 × 10−5 minocycline (pH 7.4, bubbled with 95% O_2_, 5% CO_2_). Slices were then cut using a vibrating blade microtome (Campden model 7000smz2) in an ice-cold solution containing (mM): 150 sucrose, 10 choline-Cl, 2.5 KCl, 3.1 Na-pyruvate, 11.6 ascorbic acid, 1.25 NaH_2_PO_4_, 26 NaHCO_3_, 30 glucose, 0.5 CaCl_2_, 7 MgCl_2_, supplemented with D-APV (25µM). They were then transiently immersed (10/15min) in a bath containing the same solution at 32°-34°C (pH 7.4, bubbled with 95% O_2_, 5% CO_2_). Finally, slices were transferred to warm BBS (32°-34°C) for the rest of the experimental day.

In our hands, SwiChR was tonically activated by ambient light. Throughout the experimental day, we thus paid particular attention to protect slices from all light sources.

For electrophysiology, slices were moved to a recording chamber mounted on an upright Olympus BX51WI microscope (Olympus France) and continuously perfused with bubbled BBS (∼1/1.5 ml/min; 32–35°C). Cells were visualized with a combination of Dodt contrast, and an on-line video contrast enhancement. Whole-cell (WC) recordings of PVT/CR+ cells were performed with an EPC-10 double amplifier (Heka Elektronik, Lambrecht/Pfalz, Germany) run by the PatchMaster software (Heka). PVT cells could be easily identified in the transmitted deep red light (∼750 nm) with which slices were visualized using a CoolSnap HQ2 CCD camera (Photometrics, Trenton, NJ) run by MetaMorph (Universal Imaging, Downington, PA).

Whole-cell (WC) recordings were performed with pipettes (resistance: 2/3 MΩ) filled with an intracellular solution containing (mM): 75 NMDG, 4.6 MgCl_2_, 10 HEPES, 0.5 EGTA, 4 Na_2_- ATP, 0.4 Na_2_-GTP, 1 QX-314, ∼300 mOsm and pH 7.3 with HCl. This solution maximizes the driving force for chloride ions by containing a close to symmetrical concentration with respect to the external BBS.

Spontaneous inhibitory postsynaptic currents (IPSCs) were recorded at a holding potential of -70mV in the continuous presence of the AMPA receptor blocker NBQX (5/10µM) in the bath. Only SwiChR-expressing neurons were recorded, identified by an inward current developing in response to 470nm diode pulses (Thorlabs Maisons-Laffitte, France). Light pulses were relayed to the recording chamber via the epifluorescence pathway of the microscope. During recordings the series resistance was partially compensated (max 65%), whereas liquid junction potentials were not corrected. The analysis was performed using custom-built routines in Igor (Wavemetrics, Lake Oswego, USA).

### Data analysis

#### EEG/EMG analysis

EEG and EMG data processing was carried out using Matlab. EEG signals were down sampled at 2 kHz and low-pass filtered for 50 Hz for further analysis. Power of delta (1-4 Hz) frequency band were calculated from one of the frontal screw electrodes. EMG signal was detected either directly from the EMG electrode or indirectly from one of the parietal EEG screw electrodes. For further analysis, EMG signal was downsampled at 2 kHz and bandpass filtered between 300-600 Hz. Sleep-wake states were determined using EEG and EMG signal. Wake was characterized by high EMG activity, and low delta power, while NREM sleep was characterized by low muscle tone and high delta power.

EMG ON and OFF states were defined in the following way. EMG time series were divided to 0.1 sec bins and standard deviation (SD) was calculated for each bin. Plotting a probability distribution for SD values of muscle activity for each animal, we were able to determine a value (peak of the distribution) characteristic for muscle activity in sleep. Then, using a threshold – determined for each animal (+/- [2.1-2.5] SD of baseline) – each time bin was assigned either EMG ON (1 value) or EMG OFF (0 value).^7^

#### Sleep electroencephalogram (EEG) spectral slope analysis

Periods of non-rapid eye movement (NREM) sleep EEG spectral slope was defined according to a previously published approach ^48^: the best linear fit to the equidistantly oversampled (piecewise cubic Hermite interpolation) double logarithmic plot of the average power spectral density (Welch periodogram based on 4 s Han windowing and Fast Fourier Transformation). Fitting was based on the 2–6 Hz and 18–48 Hz ranges, in order to exclude frequencies characterized by frequent upward deflections caused by NREM-sleep specific spectral peaks. The slope of the linear is known to reflect the depth of sleep (sleep intensity) ^48, 67^.

#### Horizontal activity

The horizontal movement of the animals was calculated from the video. Using a neural network-based program package called DeepLabCut ^68^, we calculated the X and Y coordinates of the animal’s head for each video frame (Fig. 2, B-lower panel, C-lower panel). DeepLabCut’s neural network was pre-trained on our own data. We then filtered the coordinates based on their likelihood (at least 0.8) and calculated the magnitude of displacement for each frame. For Fig. 1D, Fig. 3E and Fig. 8B figures we averaged 15, one-minute-long displacement data. For Fig. 1E, Fig.3 F 8D the displacement data set was normalized by the averaged nest onset time of each animal calculated for the PRE period.

#### Analysis of behavior in the nest

We defined the following behavioral variables:

##### Nest onset

Nest onset was defined as the moment when an animal enters its nest and did not leave it for at least 20 seconds.

##### Sleep onset

We defined sleep onset as the time point when the first 2 minutes of uninterrupted sleep was initiated. Sleep onset was defined using the combination of EEG and EMG (three times larger amplitude of the EEG activity compared to baseline and EMG OFF state).

##### Nest time

Nest time was defined as the time between nest onset and sleep onset.

For the detailed analysis of the behavior we used the Solomon Coder software from the beginning of the recording until the sleep onset of the animals. During the analysis we differentiated and quantified the following behaviors:

##### Nest building

A stereotyped mouse behavior ^69, 70^, during which the mouse usually reaches out, pulls in or gathers nest material to build its nest (Movie S5).

##### Freezing in the nest

During this freezing-like behavior in the nest, the animals were awake, the body of the animal did not perform horizontal movements for at least 2 s (range 2-60 s). During this type of behavior the animals sometimes perform sudden jerky movements, head-bobbing or turning but no nest building, tramping, nest tromping ^69, 70^ comfort movements or rearing (Movie S6).

##### Hyperventilation

On POST 0 day, immediately after the stress animals displayed hyperventilation visible in the video. This type of heavy breathing was only detected when the animal was at rest, as it is not visible during movement (Movie S3).

#### Unit clustering

The clustering was done in a semi-automatic way. The 3 hours of measurement were recorded in 2 files of 1.5 hours each. These files were automatically analyzed with SpyKING Circus ^71^. SpyKING Circus automatically generated clusters. These clusters were then manually curated using the Phy program ^72^. All clusters with an average amplitude below 50 <u>µ</u> V were discarded (fig. S4, A, E). Noise was then filtered out from the clusters by displaying the different PCA component features of the channels and cutting the resulting clusters (fig. S4, E). In doing so, it was checked that a given unit really belonged to a single cell and that the same cell was not scattered in several clusters (using Amplitude View and Feature View). Then, the autocorrelograms (fig. S4, C) and inter spike interval distributions of the cells were checked (no spike within 2 ms and ISI distribution with Gaussian curve shape were accepted).

#### Unit activity analysis

##### State modulation index (SMI)

All recordings were split into three behavioral stages based on the camera recordings, namely ‘Wake’, ‘Nest’ and ‘Sleep’. For sleep we get an approximately identical time window to Wake and Nest state (first 20-30 min. of sleep), this way in unit data sleep consists mainly NREM stages. The average firing rate of each neuron was computed for each stage, normalized to its time and to the average firing rate of each state on the PRE period. To quantify the effect of behavioral state on the firing of the PVT/CR+ neurons, we defined a state modulation index (SMI) as: (a-b)/ (a+b) where “a” is the proper firing rate of the PVT/CR+ neuron in state with higher activity of the comparison (i.e. wake or nest) and “b” is the proper firing rate of the cell with lower activity (i.e. nest or sleep).

##### High Frequency Activity (HFA)

HFA was defined as a series of two or more action potentials with interspike intervals less than 10 ms (fig. S7, A). Autocorrelograms were calculated at 1 ms resolution for each neuron. Normalized values were defined as the ratio of spikes in high frequency clusters within all spikes (fig. S7, C).

##### Correlated activity (CA)

CA was calculated for all simultaneously recorded pairs of neurons. Crosscorrelograms were calculated for each pair at 1 ms resolution. Correlation events between cell pairs were defined as the spikes of a neuron within ± 5 ms with a lag from the spikes of the other neuron (fig. S5, B). To characterize how likely is that pairs of neurons fire together in a ± 5 ms time window, we calculated the mean counts in the ± 5 ms lags of the CCGs and normalized these values with the mean counts in the 180-200 ms baseline period (fig. S5, D).

##### Cell tracking among days

For identification of the same cell between days we took the average action potential shape of each cell for each channel of the tetrode. To do this, we clipped the signal shape around the cell action potential peak times in a ± 2 ms window, sampled at 15 kHz from the original signal high-pass filtered at 250 Hz. Then we identified the channel with the highest peak amplitude and used this value to normalize the waveforms with the largest peak amplitude in each 4 channels of the tetrode. We quantified the waveform similarity between all recorded single unit activities by taking the sum of the squared timepoint-by-timepoint differences in the 4 channels (Fig S3, B). Finally, we examined the distribution of these points and identified the most similar 5% of the pairs as neurons recorded more than once on consecutive days. These pairs had similarity scores less than 2.75 µV∧2 (Fig S3, C). From these, we selected cells on the corresponding days (PRE 5 days and POST0 days) for each animal. In case of these cells, the similarity score ranged between 0 and 1.32 (Fig S3, B and C).

##### Optotagging with ChR2

Stimulus-Associated spike Latency Test (SALT) was used to identify significantly light-activated PVT/CR+ neurons in control animals (ChR2) ^73^. Neurons activated at p < 0.01 were considered identified CR+ neurons.

##### Optotagging with SwiChR

During SwiChR tagging, we used the protocol, detailed above in Optogenetic inhibition and phototagging section. For each cell we calculated the firing rate in the second before laser ON and in the 2^nd^ second of the laser ON period and computed the change between them using the formula (a-b)/ (a+b) was used. We measured inhibition using this ratio and considered as inhibited if this value was above 0.1 (fig. S1).

### Statistics

Data are represented as mean ± SEM. All the statistical analysis was done using GraphPad Prism version 9. First, normality of the data distribution was checked, using Shapiro-Wilk normality test. When comparing two groups, if both groups showed a normal distribution, a t-test was used. Paired t-tests were used when the same animal or cell was tested over several days (indicated by a connecting line between the 2 points in the figures), and unpaired t-tests in all other cases. If the requirements of t-test were not fulfilled, then Mann-Whitney test or Wilcoxon matched-pairs signed rank tests were used. To compare multiple groups, one-way ANOVA was used followed by Tukey’s posthoc test. The significance level was set at p *<* 0.05. Individual data points are shown in the figures. All the statistics used in figures and supplementary figures are shown in tables of statistics (**Table S1 and S2**).

