## Supplementary Figures for "Persistently increased post-stress activity of paraventricular thalamic neurons is essential for the emergence of stress-induced maladaptive behavior"

### Supplementary figures and legends

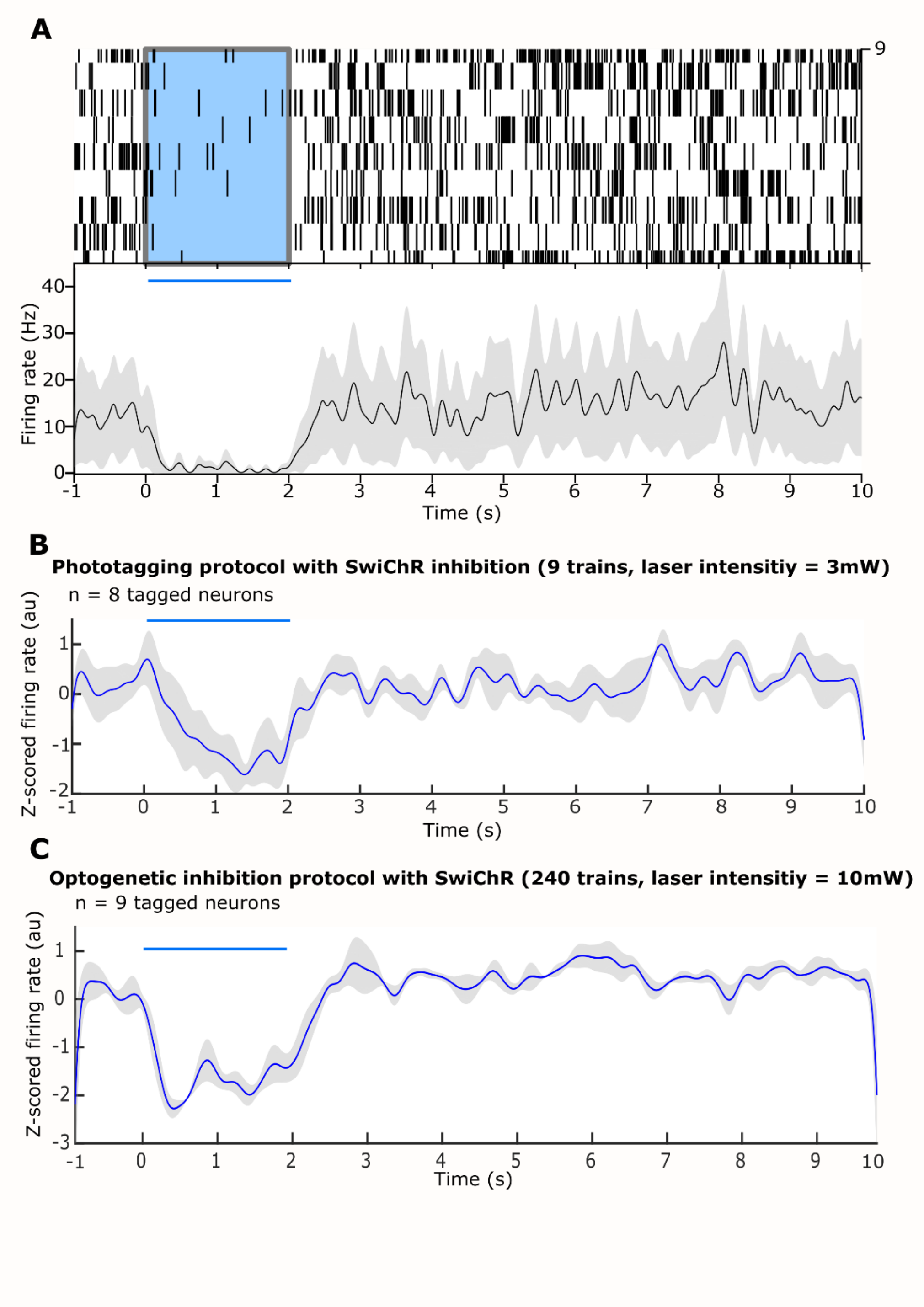


**Fig. S1. Effect of SwiChR photoinhibition on PVT/CR+ neurons during tetrode recordings.**

(A) Representative example of the firing rate of a PVT/CR+ neuron during 2s laser ON 13 sec laser OFF trains (n=9 trains) in the wake state. *Top* Raster plot of one neuron during 9 trains. Blue rectangle marks the laser ON time (2s). *Bottom* Firing rate for the same cell calculated in 10 ms windows.

(B) Z-scored firing rate of 8 phototagged PVT/CR+ neurons using the 4x15 sec phototagging protocol (see Methods). Mean (blue line) +/-SD (grey) are shown. Straight blue line above marks the laser ON time (2s).

(C) Z-scored firing rate of 9 phototagged PVT/CR+ neurons during the 60 min long photoinhibition protocol (see Methods). Mean (blue line) +/-SD (grey) are shown. Straight blue line above marks the laser ON time (2s).

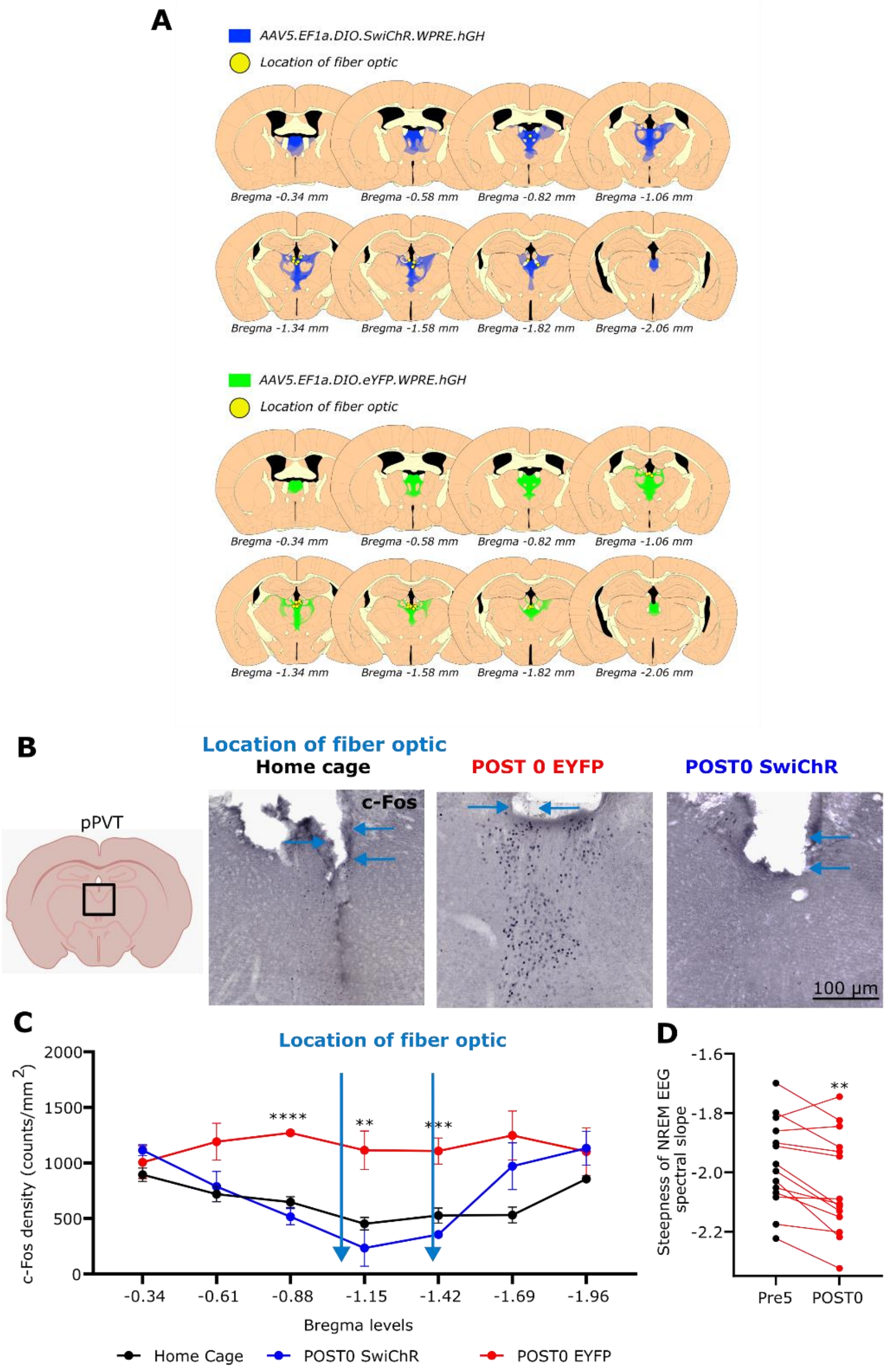


**Fig. S2. Histology of the behavioral experiments, c-Fos expression across the rostrocaudal extent of the PVT and EEG sleep slopes on POST0 days**

(A) Schematics of coronal sections illustrating the location of the optic fibers (yellow dots) and the extent of transfection following SwiChR (top, blue) and EYFP (bottom, green) virus constructs injected to the PVT of CR-Cre mice. Drawings are based on a compilation of 14 animals for EYFP and 7 for SwiChR. The schematics of coronal sections was created according to the Franklin and Paxinos mouse brain atlas (*50*).

(B) Representative images showing c-Fos immunolabeling in the PVT with fiber optic tracks (blue arrows) from home cage control, EYFP, and SwiChR mice after POSE.

(C) Quantification of c-Fos expression across the rostro-caudal extent of the PVT. Blue arrows mark the position of the fiber optics (-0.88 Bregma level, $F(2,10) = 58.33$, $p = 0.0001$; -1.15 Bregma level, $F(2,13) = 12.44$, $p = 0.001$; -1.42 Bregma level, $F(2,11) = 18.53$, $p = 0.0003$).

(D) Comparison and the steepness of the NREM EEG spectral slope between the $5^{th}$ day (black dots) of the pre-stress period and POST0 day (red dots) in EYFP mice ($t[13] = 3.58$, $p = 0.0034$). Dots represent individual animals.

See Table S1 for the full results of the statistical tests. Data are means ± SEM. *p < 0.05, **p < 0.01, ***p < 0.001, ****p < 0.0001.

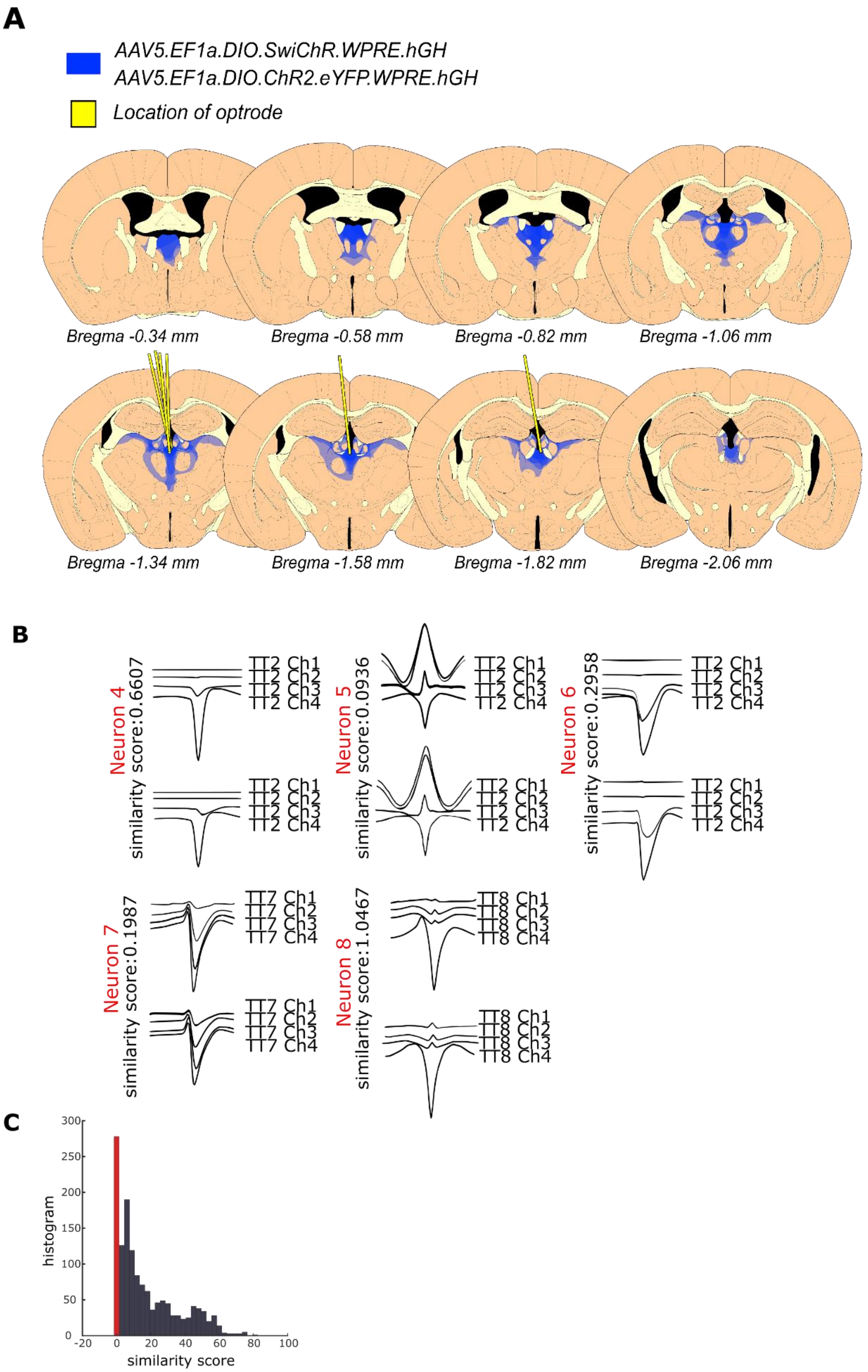
A
AAV5.EF1a.DIO.SwiChR.WPRE.hGH
AAV5.EF1a.DIO.ChR2.eYFP.WPRE.hGH
Location of optrode
Bregma -0.34 mm
Bregma -0.58 mm
Bregma -0.82 mm
Bregma -1.06 mm
Bregma -1.34 mm
Bregma -1.58 mm
Bregma -1.82 mm
Bregma -2.06 mm
B
Neuron 4
similarity score:0.6607
Neuron 5
similarity score:0.0936
Neuron 6
similarity score:0.2958
Neuron 7
similarity score:0.1987
Neuron 8
similarity score:1.0467
TT2 Ch1
TT2 Ch2
TT2 Ch3
TT2 Ch4
TT7 Ch1
TT7 Ch2
TT7 Ch3
TT7 Ch4
TT8 Ch1
TT8 Ch2
TT8 Ch3
TT8 Ch4
C
histogram
similarity score
0
50
100
150
200
250
300
-20
20
40
60
80
100

**Fig. S3: Histological verifications of the tetrode experiments and tracked neurons from PRE5 day to POST0 day in ChR2 animals**

(A) Schematic of coronal sections illustrating the placement of tetrodes (yellow lines) and extent of injection sites using conditional ChR2 or SwiChR (blue) containing virus constructs targeted to the PVT in CR-Cre mice. Drawings are based on 6 mice (n=4 for ChR2 and n=2 for SwiChR). The schematic of coronal sections was created according to the Franklin and Paxinos mouse brain atlas (*50*).

(B) Representative waveforms of optotagged PVT/CR+ neurons in different tetrodes of the same animal shown in Fig4E (TT2, TT7, TT8). The neurons were recorded for two consecutive days (top vs bottom row). Channel numbers (Ch) and similarity scores between the two days are shown next to the waveforms.

(C) Distribution of similarity scores of all tagged PVT/CR+ neurons from ChR2 and SwiChR animals. Red line indicates the threshold.

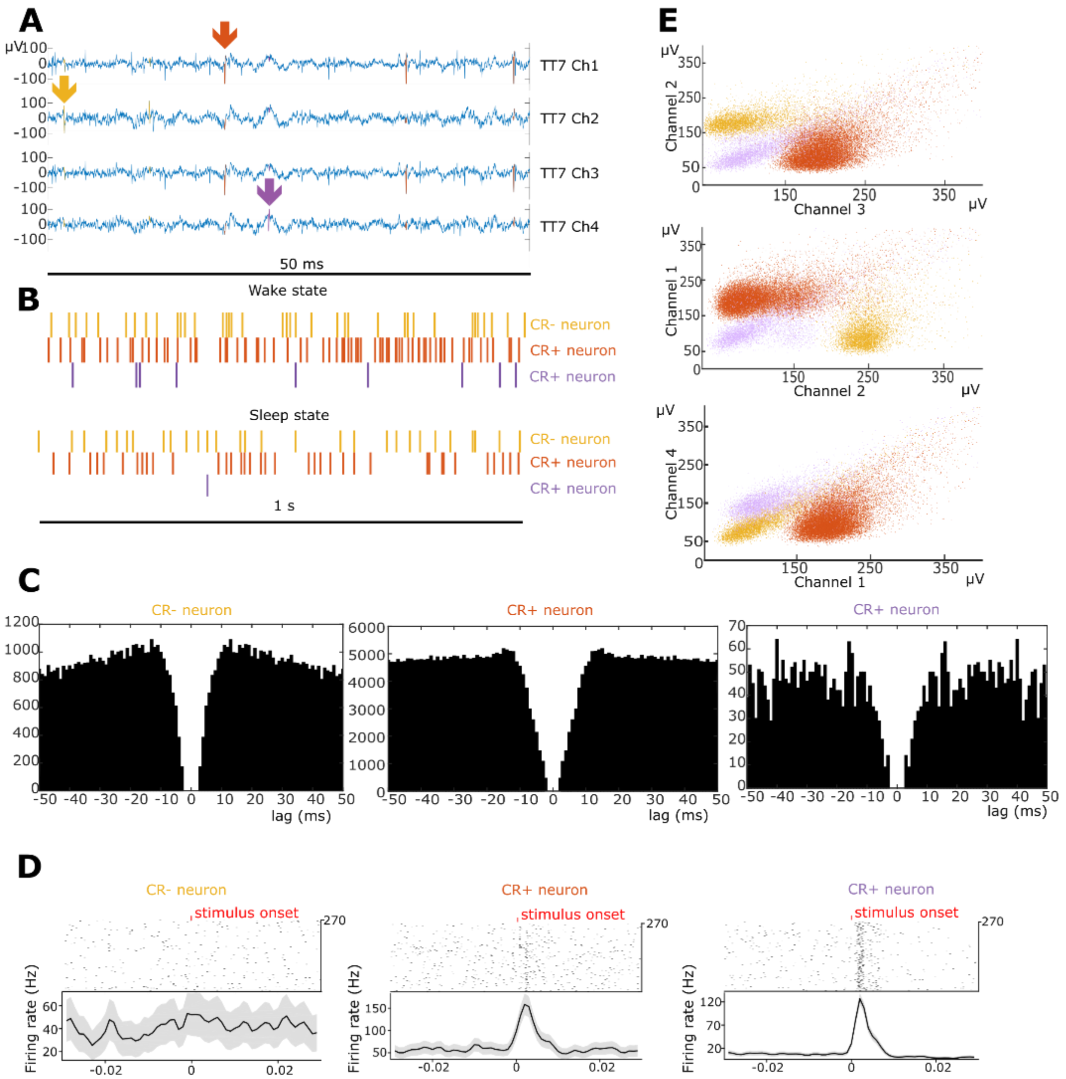


**Fig. S4: Representative example of the clustering process in case of three individual neurons recorded on the same tetrode in the PVT.**

(A) Action potentials (arrows, colored by yellow, orange and purple) of three PVT units identified on a 50 ms long raw trace (high pass filtered, above 150 Hz).

(B) Timing of action potentials of the same three neurons on a 1 s long window (two of them is CR+ one of them are CR-).

(C) Autocorrelograms of these three PVT neurons, calculated from the 3 hours long recording with 1 ms bins in a 50 ms window.

(D) Peristimulus time histogram of the optogenetic stimulation (tagging) of the same three neurons during wakefulness in the 30 ms before and after the stimulus. Red ticks mark the onset of the 1 ms long stimulation. One non-tagged (yellow, putative CR-) and two tagged (orange and purple, CR+) neurons are shown.

(E) The three amplitude clusters of the same three neurons in the three channels used for clustering (one of the PCA components during clustering).

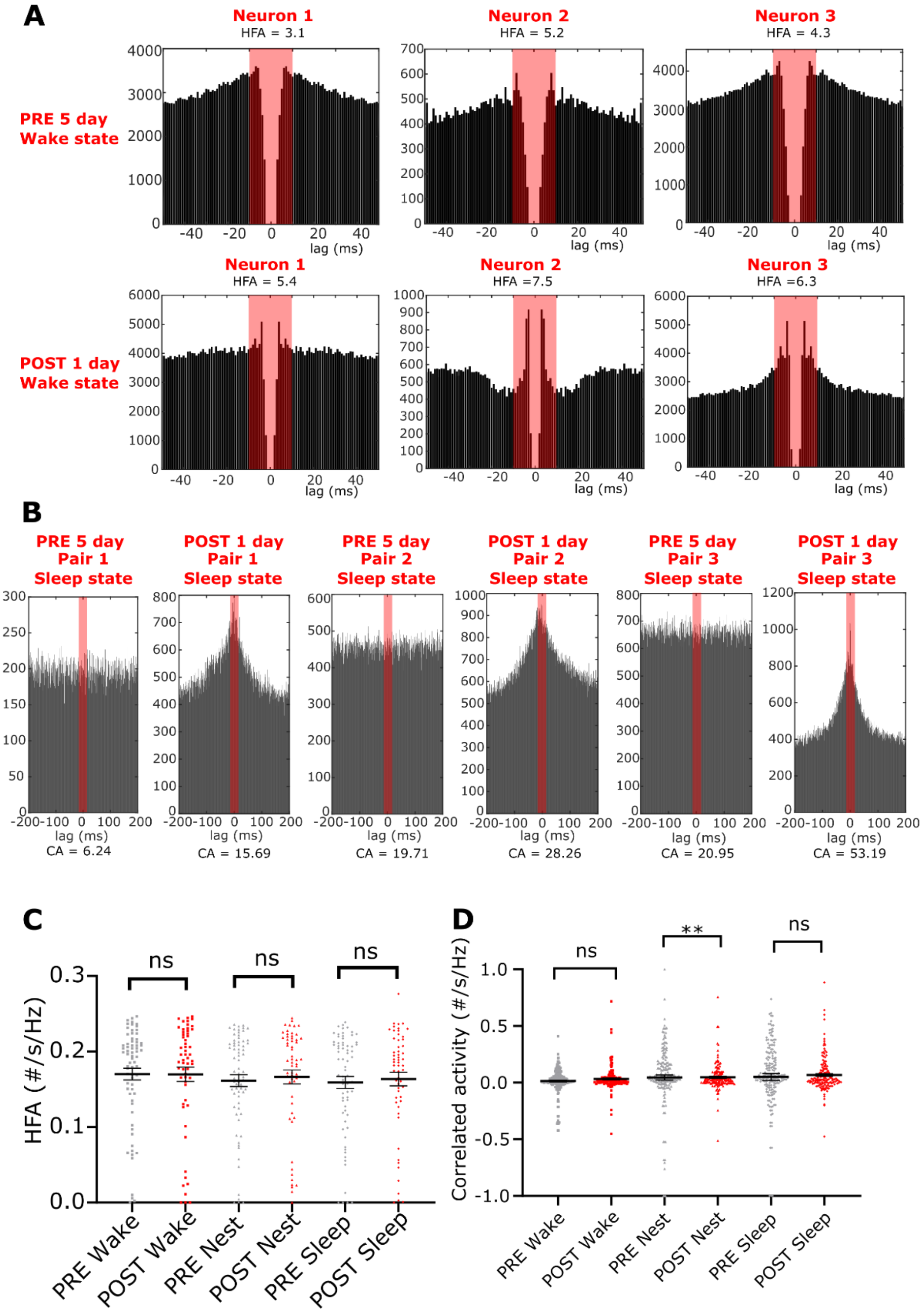


**Fig. S5. Representative examples and normalized values of High Frequency Activity (HFA) and Correlated Activity (CA) in ChR2 tagged neurons.**

(A) Autocollerograms of three tracked neurons with 50 ms lag recorded on the PRE 5 day (before the stress) and on the POST1 day (after the stress) during wake state. The red shading shows the time window (-10;10 ms) used to compute HFA values (shown above the autocorrelograms). Note pronounced change in HFA activity (large central peaks) in Neuron 2 after the stress.

(B) Crosscorrelograms of three tracked pairs of neurons with 200 ms lag on the PRE 5 day (before the stress) and on the POST1 day (after the stress). The red shading shows the time window within (-5;5 ms) used to compute CA values (shown below the crosscorrelograms). CA values increase in all three pairs after the stress.

(C) HFA values normalized with mean firing rate (see Methods) of the recorded and tagged neurons from control (ChR2) animals on the PRE (n = 69 from 4 animals) and the POST (n = 57 neurons from 4 animals) period (wake, U = 1912, p = 0.792; nest, U = 1786, p = 0.3789; sleep U = 1910, p = 0.7845).

(D) CA values normalized with baseline activity (see Methods) of the recorded and tagged neurons from control (ChR2) animals on the PRE (n = 69 from 4 animals) and the POST (n = 57 neurons from 4 animals) period (U = 15337, p = 0.8632; nest, U = 12969, p = 0.0082; sleep U = 14073, p = 0.1357). See Table S1 for the full results of the statistical tests. Data are means ± SEM. **p < 0.01.

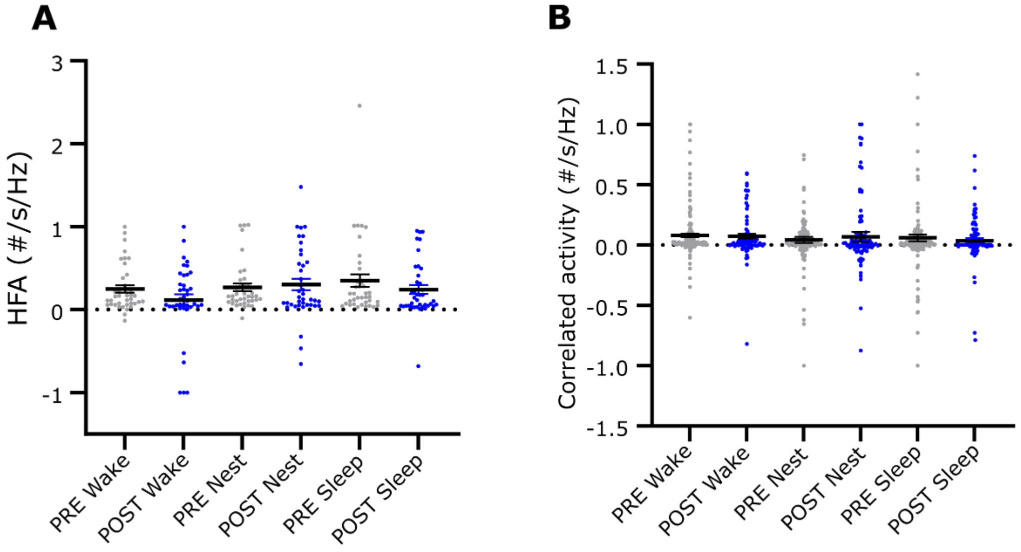


**Fig. S6. Normalized values of High-Frequency Activity (HFA) and Correlated Activity (CA) in SwiChR tagged neurons.**

(A) Normalized HFA values with firing rate (see Methods) of the recorded and tagged neurons from inhibited (SwiChR) animals on the PRE (n = 38 from 2 animals) and the POST (n = 39 neurons from 2 animals) period (wake, U = 870.5, p = 0.5301; nest, U = 891, p = 0.6496; sleep U = 862, p = 0.4854).

(B) Normalized CA values with baseline activity (see Methods) of the recorded and tagged neurons from control (SwiChR) animals on the PRE (n = 38 from 2 animals) and the POST (n = 39 neurons from 2 animals) period (U = 7306, p = 0.552; nest, U = 7418, p = 0.6884; sleep U = 7356, p = 0.6106). See Table S1 for the full results of the statistical tests. Data are means ± SEM.

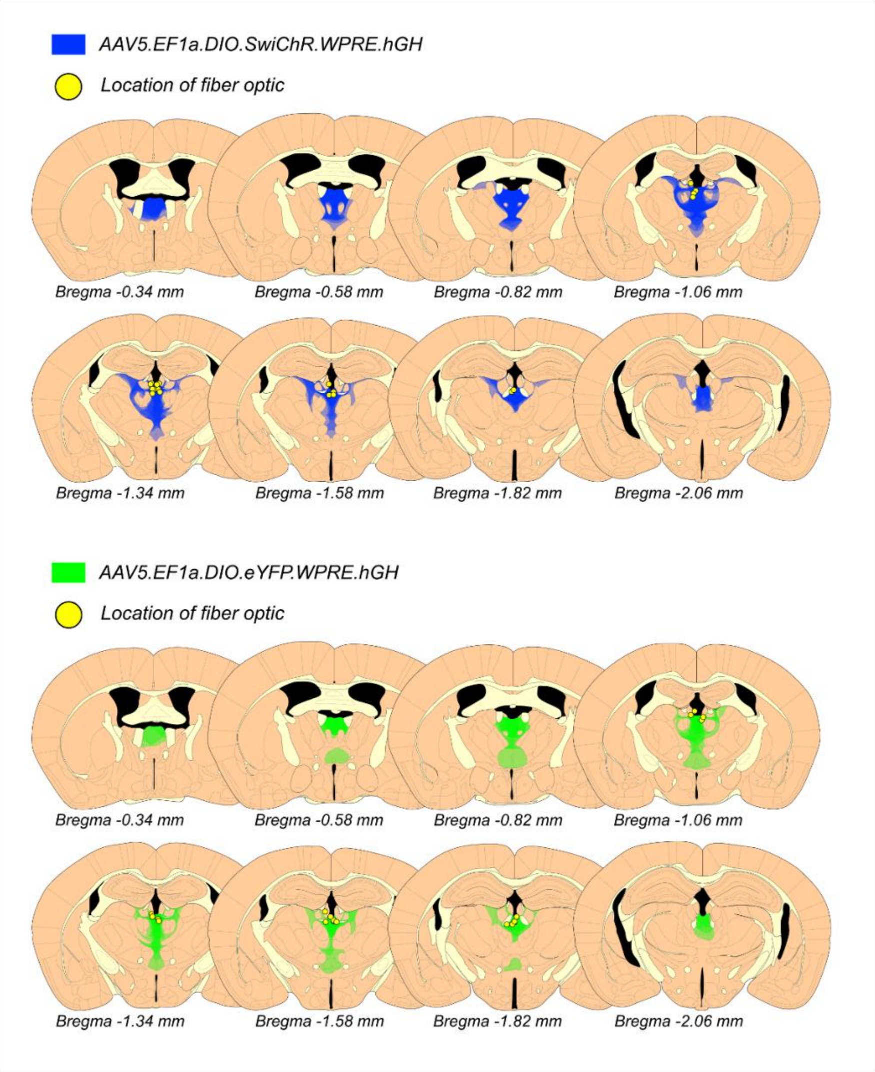


**Fig. S7. Injection sites and fiber optic locations of mice involved in GABA A receptor and c-Fos mapping experiments.**

Schematics of coronal sections illustrating the location of the optic fibers (yellow dots) and the extent of transfection following SwiChR (top, blue) and EYFP (bottom, green) virus constructs injected to the PVT of CR-Cre mice. Drawings are based on a compilation of 10 animals for EYFP and 10 for SwiChR. The schematics of coronal sections was created according to the Franklin and Paxinos mouse brain atlas (*50*).

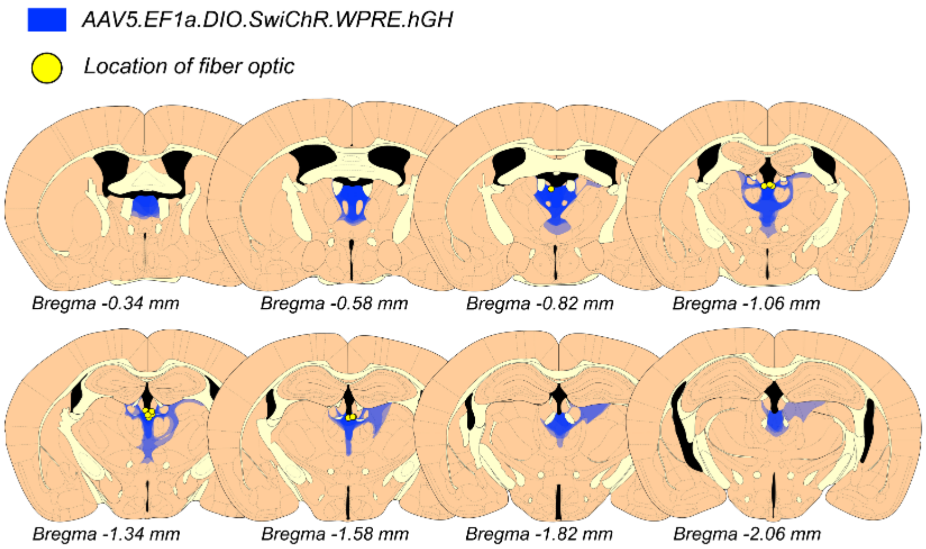


**Fig. S8. Injection sites and fiber optic locations of mice involved in LATE photoinhibition experiment.**

Schematics of coronal sections illustrating the location of the optic fibers (yellow dots) and the extent of transfection following SwiChR virus constructs injected to the PVT of CR-Cre mice. Drawings are based on a compilation of n=7 SwiChR mice. The schematics of coronal sections was created according to the Franklin and Paxinos mouse brain atlas (*50*).

### SUPPLEMENTARY MOVIES LEGEND

**Movie S1**. Two representative animals from the EYFP group during the predator odor stress exposure (POSE) protocol displaying strong defensive behavior, mostly freezing and escape jumps. The capsules containing 2MT are located in lower part of the cage. POSE is performed in a cage distinct from the home cage in a different room, under a fume hood.

**Movie S2.** Two representative animals from the no odor exposure (NOE) group in the same environment. Animals display normal explorative behavior.

**Movie S3.** Behavior of an animal from the EYFP group displaying hyperventilation and freezing, immediately after POSE when the animal was returned to its home cage.

**Movie S4.** Behavior of an animal from the SwiChR group displaying normal locomotor behavior, immediately after POSE when the animal was returned to its home cage. The 2 sec light pulse on the head implant indicates the laser ON periods.

**Movie S5.** Characteristic nest building behavior of an animal from the EYFP group during the PRE period. The animal collects, processes, and arranges the paper used as nesting material.

**Movie S6.** Characteristic behavior of an animal from the EYFP group during the POST period. The animal mainly displays freezing-like behavior (shortly freezing) in the nest. During this type of behavior, the animal is largely stationary, does not perform horizontal movements (for at least 2 s) just sudden jerky movements, head-bobbing or turning. Nest building or rearing are rare.
